# Direction-Dependent Encoding of Forward and Inverse Models in the Same Purkinje Cells

**DOI:** 10.64898/2026.11.02.763589

**Authors:** Junya Inoue, Akshay Markanday, Peter Thier

## Abstract

Saccadic eye movements are too fast to be guided by sensory feedback and therefore depend on precisely calibrated internal models. The cerebellum is widely thought to compute both forward and inverse models, yet how these two computations coexist in the same circuit remains unclear. We hypothesized that the same Purkinje cells (PCs) can serve as a forward model for saccades in one direction and as an inverse model for saccades in the opposite direction, because target neurons on either side of the cerebellar nuclei contribute differently to leftward and rightward saccades. To test this hypothesis, we recorded from PCs in the oculomotor vermis of macaque monkeys during repetitive saccade paradigms that induce adaptation and fatigue. Using the preferred error direction of each PC conveyed by increased complex spike (CS) activity (CS-on direction) as a proxy of the side of the cerebellar nuclei it projects to, we compared population activity for saccades directed against the CS-on direction (CS-on opposed) and along it (CS-on aligned). CS-on opposed saccades, (toward the PC’s projection side) were accompanied by a precisely timed, sharp burst of simple spike (SS) activity at saccade onset that scaled with peak velocity, and CS error sensitivity was highest after the saccade. SS activity also changed systematically after adaptation, reflecting the altered kinematics of the adapted saccade. These properties are consistent with a forward model that estimates ongoing movement and updates itself based on post-movement error. In contrast, the CS-on aligned saccades, (away from the PC’s projection side) were accompanied by broader SS modulation around saccade offset, and the CS error sensitivity was highest before the saccade. CSs around saccade onset encoded target location and initial eye position, and their occurrence was associated with reduced saccade amplitude. This reduction likely adjusts the motor command to the starting eye position, as expected for an inverse model that pre-programs the motor command before movement begins. Together, these findings provide neural evidence that the same PC population can contribute to both forward and inverse model computations, depending on saccade direction.

## Introduction

Saccades are rapid, ballistic eye movements, yet they land on the target of interest with remarkable precision. As saccades are too brief for visual feedback to guide them, and visual sensitivity is reduced around the time of saccades (Matin, 1974; Volkmann, 1986), saccadic control must rely on precisely calibrated internal models. In motor control, there are generally two types of internal models, forward models and inverse models (Wolpert et al., 1998). In forward models, an efference copy of the motor command is used to predict the expected movement kinematics, and this prediction is then used to update the ongoing motor command. Because the sensory prediction is available with minimal delay it can substitute for veridical sensory feedback within the control loop, which is often too slow to support online correction. An inverse model, on the other hand, computes the motor command needed to realize the desired movement kinematics based on the current state of the body and the environment (i.e., movement dynamics). As this computation is needed before movement initiation, this model is implemented in a feedforward pathway (Fig. 1 a).

**Figure 1:**
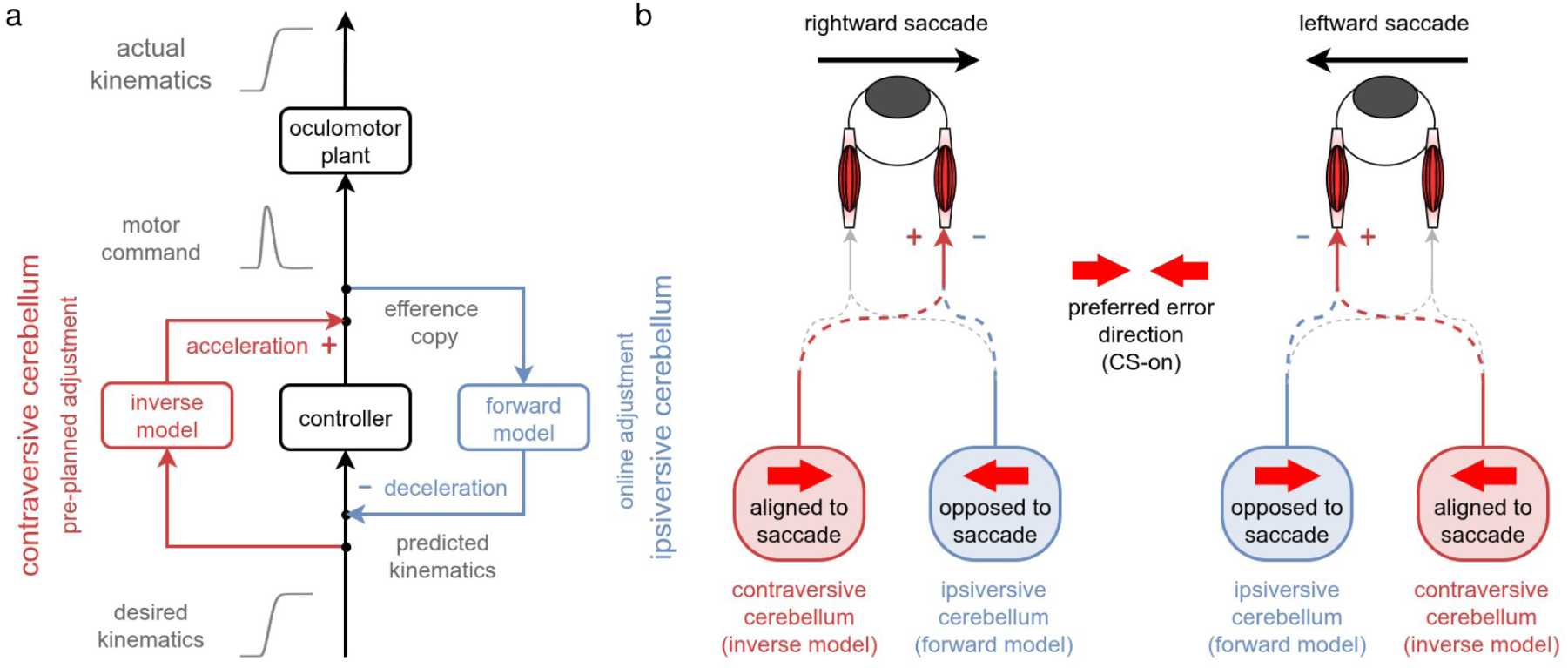
Theoretical framework. **a**: Schematic of saccade control circuitry. The inverse model computes the adjustment of the motor command required to achieve the desired movement kinematics. It is proposed to operate in a feedforward pathway in the contraversive cerebellum, where it adds a pre-planned drive that accelerates the eye (+) based on the desired kinematics. This adjustment must therefore be specified before movement onset. The forward model predicts the kinematics resulting from the issued motor command. It is proposed to operate in a feedback loop in the ipsiversive cerebellum, where it uses an efference copy of the motor command to predict the ongoing movement kinematics, which allows the command to be adjusted online to decelerates the eye and stop it at the right time (-). The controller corresponds to premotor burst neurons in the brainstem. **b**: Direction-dependent roles of the same cerebellar side. Left: for a rightward saccade, the cerebellar side whose PCs have a rightward CS-on direction (red, aligned to the saccade) is contraversive and acts as an inverse model, where as the side with a leftward CS-on direction (blue, opposed to the saccade) is ipsiversive and acts as a forward model. Right: for a leftward saccade, the roles are reversed, so each side switches between the inverse and forward model depending on saccade direction. Red and blue arrows indicate excitatory (+) and inhibitory (-) influences on the agonist eye muscle, and gray dashed arrows indicate pathways not driving this muscle. Red block arrows indicate the preferred error direction (CS-on) of each cerebellar side. Contraversive and ipsiversive refer to the cerebellar side relative to the saccade direction. Cerebellar output reaches the eye muscles via brainstem saccade generator (not shown).

The cerebellum is a strong candidate to compute the internal models serving movement control. This is suggested by a large body of findings on its architecture (Albus, 1971; Marr, 1969; Wolpert et al., 1998), the properties of cerebellar neurons (Itō, 1984; Shidara et al., 1993) as well as the consequences of cerebellar lesions (Barash et al., 1999; Bastian et al., 1996; Gerwig et al., 2003; Ignashchenkova et al., 2009). For instance, cerebellar impairment leads to characteristic disturbances of saccades. First, it causes saccadic dysmetria, i.e., saccades that systematically miss the target by overshooting or undershooting it, or by deviating from its direction. Second, it impairs saccadic adaptation, i.e., the adjustment of saccade gain in response to movement errors. Third, although saccades may on average be on target, the variability of saccadic endpoints is typically increased (Barash et al., 1999). Lastly, the saccades become slowed with reduced peak velocity (PV) and disproportionately prolonged duration (Bürk et al., 1999; Buttner et al., 1998; Golla et al., 2008; Robinson et al., 1993). These saccadic abnormalities caused by cerebellar impairments can be explained by inaccurate state estimates in the forward model or erroneous motor command in the inverse model. Specifically, a disrupted forward model would fail to accurately predict the current eye state, leading to dysmetria, increased endpoint variability, and inability to adapt to novel systematic errors. A miscalibrated inverse model, on the other hand, would generate inappropriate motor commands from the outset, which could produce dysmetria as well as abnormal saccade kinematics such as reduced PV and prolonged duration. The cerebellum is well-suited to accommodate the computations of both types of internal models serving saccades as well as other types of sensory-guided movements as it has access to motor commands and sensory state estimates of the eyes, the rest of the body and the environment conveyed by afferents from the cerebral cortex, brainstem, and spinal cord. To understand how these computations may be implemented at the circuit level, we briefly outline the key architectural features below.

Cerebellar cortex is three-layered, consisting of an innermost layer of granule cells (GCs), a middle layer of Purkinje cells (PCs) and an outermost molecular layer comprising axonal and dendritic fibers with interspersed molecular layer interneurons (MLIs). GCs receive information from various area of the central nervous system, including the cerebral cortex, brainstem nuclei and spinal cord, via mossy fibers (MFs) (Itō, 1984). Their axons give rise to parallel fibers (PFs), and each PC receives more than 100,000 PF synapses (Napper and Harvey, 1988). These PF-PC synapses are widely believed to be the primary site of internal model computations, as they provide the synaptic capacity to store and update the mappings between motor commands and their sensory consequences. PCs convey the results of these computations via simple spikes (SS), driven by MF-PF input and intrinsic activity, to the deep cerebellar nuclei (DCN),which relay them to downstream stages of motor control. PCs serving saccades reside primarily in vermal lobules VIc and VIIA, the oculomotor vermis (OMV) (Noda and Fujikado, 1987). Their saccade-related SS patterns are highly heterogeneous, including increases, decreases, and biphasic or multiphasic discharge, which has made it difficult to establish how SS activity might convey internal-model-like information. The second afferent system, the climbing fibers, originates from the inferior olive (IO). Each PC is contacted by only one or a few climbing fibers, which evoke complex spikes (CSs), low-frequency spikes of long duration and complex shape that serve as error signals guiding motor learning. CS affect SS output in two says. First, a CS coinciding with PF activation induces long-term depression (LTD) of the co-activated PF-PC synapses (Ito and Kano, 1982), reducing SS firing. Second, as a consequence of their impact on voltage- and Ca *^2^*^+^-dependent ion channels and the transient activation of MLIs (Mathews et al., 2012), CSs are followed by a pause in SS firing of about 20 ms (Bloedel and Roberts, 1971; Davie et al., 2008; Markanday et al., 2020). Because PCs are inhibitory, this pause disinhibits DCN neurons and may contribute to fine adjustments of ongoing movements.

The OMV projects largely ipsilaterally to the caudal fastigial nuclei (cFN), the most medial pair of DCN, so each cFN neuron receives input from PCs of the same side. Unilateral muscimol inactivation of the cFN produces direction-specific dysmetria: after inactivation of the left cFN, ipsiversive (leftward) saccades overshoot and contraversive (rightward) saccades undershoot (Robinson et al., 1993), where ipsiversive and contraversive refer to saccades directed toward or away from the inactivated nucleus. This suggests that the contraversive cFN accelerates the saccade whereas the ipsiversive cFN decelerates it. Optogenetic perturbation of OMV PCs during saccades has recently led to the proposal that these two roles might reflect the two internal models: an inverse model acting through the contraversive side to generate the accelerating drive, and a forward model acting through the ipsiversive side to decelerate the eye (Soetedjo and Horwitz, 2024). This proposal rests on the behavioral consequences of the perturbation, whereas the neural implementation remains unknown, namely whether the activity of PCs themselves shows such a direction-dependent dissociation. If this is correct, then because contraversive and ipsiversive are defined relative to the side of the cerebellum, the same PC should carry inverse-model-like signals for saccades in one direction and forward-model-like signals for saccades in the opposite direction (Fig. 1 b). Individual OMV PCs would thus be components of both forward and inverse models, depending on movement direction.

Testing this prediction requires knowing which side of the cFN a recorded PC projects to. Inferring this from the location of the recording site alone is unreliable as the vermis is a relatively small midline structure without a clear separation between its left and right sides. However, the cerebellum is organized into microcircuits, each consisting of PCs that receive climbing fiber input from a common region of the IO and project to a common target in the DCN, here the cFN (the IO-PC-cFN microcircuit). Each microcircuit serves as a functional unit of the cerebellum. Because climbing fibers signal sensory error, PCs of the same microcircuit share a CS error tuning: the CS-on direction, defined as the direction of sensory error that elicits the highest CS rates(Soetedjo et al., 2008). The CS-on direction therefore identifies the microcircuit a PC belongs to and, through it, the cFN it targets. This is consistent with the spatial organization of the OMV, where leftward CS-on PCs are predominantly found in the right OMV, and vice versa (Herzfeld et al., 2015; Pi et al., 2024). Saccades aligned with a PC’s CS-on direction are thus contraversive, and saccades opposed to it are ipsiversive, relative to its cFN target (Fig. 1 b). Here, we test our prediction at the neural level by analyzing SS and CS population activity recorded from OMV PCs in macaque monkeys, grouping neurons according to whether their CS-on direction is aligned with or opposed to the saccade direction.

## Methods

### Behavioral paradigm

Macaque monkeys were trained to perform two visually guided saccadic tasks: an *Adaptation* and a *Fatigue* task. In both tasks, a peripheral visual target appeared immediately after the central fixation point disappeared. When the monkeys made a successful saccade to the peripheral target (centrifugal saccades, CF), they received a drop of water or juice, depending on their preference, as reward. The return saccade back to the central fixation point (centrifugal saccades, CP) was not rewarded.

### Adaptation paradigm

We used an adaptation paradigm that extended and modified the paradigm originally suggested by McLaughlin (1967). It consisted of three phases: baseline, adaptation, and generalization phase (Fig. 2 a). During the *baseline phase*, a target appeared randomly at one of seven possible locations, with a fixed horizontal eccentricity of +15° (right) or - 15° (left) kept consistent within each session, and vertical eccentricities ranging from -9° to 9° in 3° increments. In the subsequent *adaptation phase*, the target appeared only along the horizontal axis, at the same horizontal eccentricity (+15° or -15°, depending on the session) as in the baseline phase. As soon as a saccade toward the target was detected, the target disappeared and reappeared, either 5° farther out or closer to the fixation point on the same horizontal axis, with the shift direction (outward or inward) fixed within a session. These intra-saccadic target shifts, typically unnoticed due to saccadic suppression, create a deceptive visual error when the eyes land at the original target location. This elicits a requirement to correct this error by making a corrective saccade to meet the target in its final, shifted position. However, the corrective saccade is not the only consequence of having initially missed the target. Rather, the amplitude of the subsequent saccades to the same initial target location is gradually increased (in outward adaptation) or decreased (in inward adaptation), depending on whether the intra-saccadic target shifts occured consistently outward or inward, respectively, to reduce the anticipated error. Finally, in the *generalization phase*, the target was again presented at one of the seven baseline locations. To maintain the adapted state and to prevent its decay, adaptation trials (those associated with intra-saccadic target shifts) were interspersed, with their proportion among horizontal-axis trials starting from 80% and gradually decreasing to 60%. Because adaptation trials were added on top of the baseline horizontal trial count, the number of horizontal-axis trials was substantially larger than the number of diagonal trials in the generalization phase (Suppl. Fig. 1). Behavioral results from a typical session are shown in Fig. 2 c-f.

**Figure 2:**
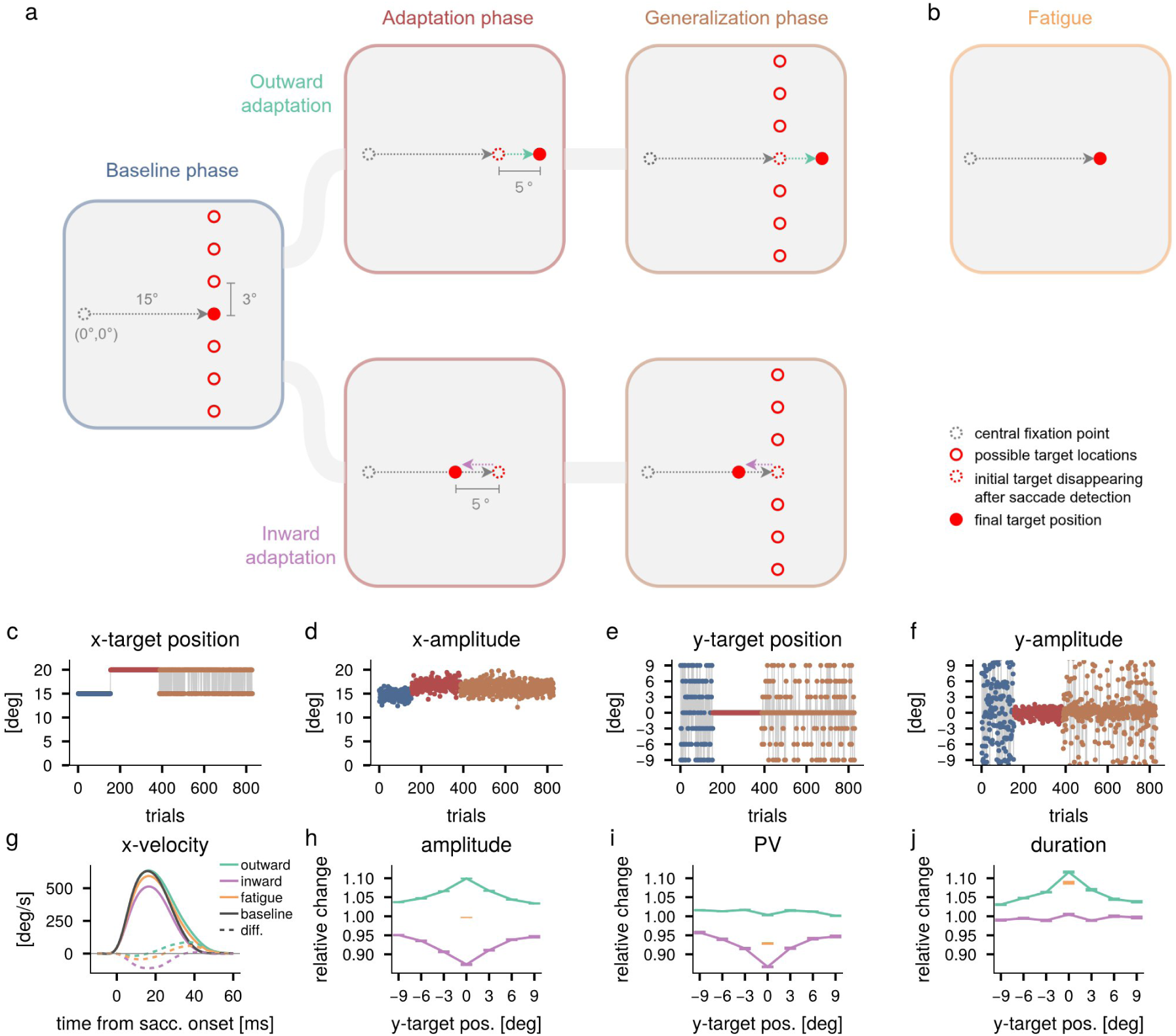
Experimental paradigms and behavioral results. **a**: Adaptation paradigm consisting of baseline, adaptation and generalization phases. During the adaptation and generalization phases, the target is shifted by 5° (outward, green arrow; inward, purple arrow) as soon as the saccade onset is detected, inducing a post-saccadic error that drives gradual updating of saccade amplitude. Targets are at 15° horizontal eccentricity ( session with +15° shown; mirrored for -15° sessions) with seven vertical positions spaced 3° apart. **b**: Fatigue paradigm. Saccadic fatigue was induced by repetition of saccades to a target always appearing in the same position (15° horizontal). **c-f**: Behavioral results from an example outward-adaptation session. Blue, red and brown dots indicate the baseline, adaptation, and generalization phase, respectively. **c**: horizontal target positions. **d**: horizontal saccade amplitude. **e**: vertical target position. **f**: vertical saccade amplitude. **g**: Mean velocity profiles during baseline phase („baseline”, gray), adaptation phase with outward shifts (“outward”, green), adaptation phase with inward shifts (“inward”, purple), and the last 50 trials of sessions from the fatigue paradigm (orange). The dashed lines show the difference between each profile and the baseline, in the color of the corresponding condition. **h-j**: Gain change of saccade kinematics in the generalization phase relative to baseline for adaptation paradigm (green: outward; purple: inward) and in the last 50 trials relative to the first 50 trials for fatigue paradigm (orange). Error bars indicate the standard error of the mean. **h**: saccade amplitude. **i**: saccadic peak velocity (PV), **j**: saccade duration.

### Fatigue paradigm

Because adaptation requires many repetitive saccades, which inevitably induce fatigue, we also ran a fatigue paradigm as a control, to dissociate adaptation-related changes from those caused by repetition alone. In this paradigm, cognitive fatigue was induced by repetitively presenting the target at the same horizontal eccentricity of 15° (no vertical offset) either leftwards or rightwards (depending on the session), requiring monkeys to execute several hundred consecutive saccades to this same location. The dataset used here is identical to the one reported in our previous studies(Markanday et al., 2023, 2021).

Unless otherwise specified, analysis of the fatigue paradigm included all trials from the entire session, combining both CF and CP saccades. The comparison between the first and last 50 trials, used to assess the effects of fatigue, is described separately where applicable.

## Results

### Direction-dependent kinematic changes after adaptation

In the adaptation paradigm, repetitive intra-saccadic target shifts led monkeys to successfully adjust their saccade amplitudes. However, the underlying kinematic changes differed between outward and inward adaptation. During outward adaptation, amplitude increase was primarily achieved through a prolongation of saccade duration because deceleration in the latter half of the saccade was weaker compared to baseline trials (green lines; the dashed line shows the difference from baseline; Fig. 2 g). In contrast, for inward adaptation, the reduction of amplitude was driven by a decrease in PV, with velocity profile shape preserved but uniformly scaled down throughout the movement (purple lines; Fig. 2 g), whereas duration did not change from the baseline trials. Notably, inward adaptation produced significantly larger relative amplitude changes than outward adaptation (horizontal saccades, outward: 0.10; inward: 0.13; Welch’s t-test; p=9.81e-8). This is not surprising, considering the influence of fatigue caused by excessive trial repetition which causes a gradual PV decline, compensated by increased duration to preserve saccade accuracy. To asses these effects, we compared PV, duration and amplitude between the first and last 50 trials of the fatigue paradigm. Consistent with previous findings (Markanday et al., 2021), fatigue reduced PV and prolonged duration without affecting amplitude (Fig. 2 h-j). During outward adaptation, fatigue counteracts the PV increase required for larger saccades, resulting primarily in prolonged duration but not PV, whereas the same fatigue facilitates the PV reduction warranted by inward adaptation, with no change in duration.

In the *generalization phase*, the amplitude changes induced by horizontal axis adaptation transferred to saccades made in diagonal directions, where no intra-saccadic target shifts were induced. The effect of adaptation decreased exponentially with distance from the trained horizontal adaptation direction, i.e. with increasing vertical eccentricity of the target. We then tested whether the decay of the adapted gains differed between the outward and inward adaptation. Given their symmetrical exponential decline in both adaptation directions, we fitted a Laplace distribution 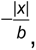 where *a* is a scaling constant and *b* reflects the decay rate, and compared *b* for the two adaptation directions. We found no statistical difference (outward: 8.04 °, inward: 9.48 °; bootstrap resampling; p = 0.10, 95% bootstrap null distribution: [-0.25, 2.58]°). Correspondingly, the key kinematic parameters underpinning the two directions of adaptation (duration for outward adaptation and PV for inward adaptation) declined equally with increasing deviation from the adaptation direction (Fig. 2 h-j).

### PC-SS population response is determined by saccade alignment relative to CS-on direction

As described in the introduction, PCs in the OMV are spatially organized according to their CS-on direction, with leftward CS-on PCs predominantly found in the right OMV and vice versa (Fig. 1 b). The CS-on direction of individual PCs can therefore serve as a proxy for their anatomical location within the OMV. To determine the CS-on direction of each PC, we compared CS firing rates for leftward and rightward post-saccadic errors during the period between 50 and 250 ms after saccade offset, assigning the direction yielding the higher firing rate as the CS-on direction. This time window was chosen consistent with our previous work (Markanday et al. (2021); see Supplementary Methods for details). Rather than using finer directional categories, we classified the CS-on direction along a single left-right axis, as our primary interest was whether the CS-on direction was aligned with or opposed to the direction of each saccade. To quantify the spatial distribution of PCs with leftward and rightward CS-on directions, we fitted a logistic regression to the probability of finding leftward CS-on PCs as a function of lateral recording position: 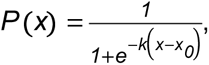 where *x* is the lateral position in recording chamber coordinates (positive to the right), *k* is the slope and *x_0_* is the midpoint of the sigmoid (both CS-on directions are equally likely *P* (*x_0_*) = *0*.*5*). This yielded *k* = *0*.*41* ± *0*.*19* and *x_0_* = *0*.*46* ± *0*.*10* [mm] (mean ±SD); leftward CS-on PCs were more prevalent in the right OMV and vice versa in our sample (Fig. 3 a), confirming that our recordings reflected the expected spatial organization (Herzfeld et al., 2015; Pi et al., 2024).

**Figure 3:**
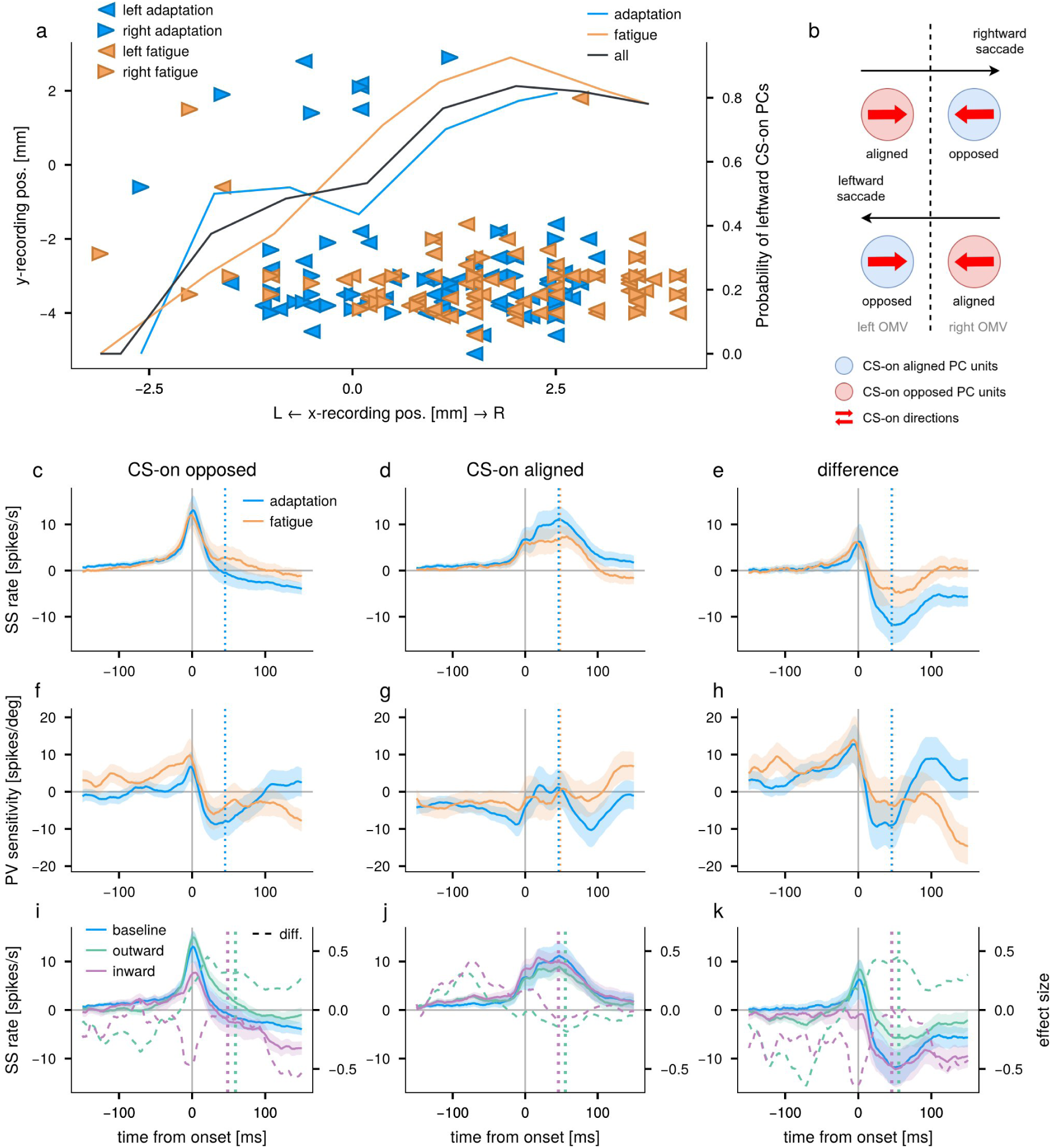
**a**: Spatial distribution of PCs plotted in the recording chamber coordinate system. Leftward- and rightward-pointing triangles indicate PCs with leftward and rightward CS-on directions, respectively. Blue and orange symbols denote PCs recorded in the adaptation and fatigue paradigm, respectively. The probability of finding PCs with a leftward CS-on direction is overlaid, with the line indicating the probability computed by combining data from both paradigms. **b**. Schematic illustrating how PCs were divided into two subpopulations based on the relationship between saccade direction and each cell’s CS-on direction. For example, for a PC with a rightward CS-on direction, a rightward saccade is a CS-on aligned while a leftward saccade is a CS-on opposed. **c**-**e**. Mean SS population response aligned to saccade onset for CS-on opposed saccades (c), CS- on aligned saccades (d), and their differences (e). SS responses from the adaptation and fatigue paradigms are shown in blue and orange, respectively. Shaded areas indicate the standard error. Dashed vertical lines indicate the mean saccade offset, colored by paradigm. **f**-**h**: Same as c-e, but for SS sensitivity to PV. **i**-**k**: Same as c-e but showing adaptation-induced changes in SS population firing rate. The blue baseline traces are identical to those shown in c-e. Solid green and purple lines depict SS responses during the outward and inward adaptation phase, respectively. The dashed lines show the effect size of the deviation from the baseline. Dotted vertical lines mark the mean saccade offset, colored by adaptation direction.

Next, we separated SS responses based on whether saccade direction was aligned with or opposite to the PC’s CS-on direction, hereafter referred to as *CS-on aligned* and *CS-on opposed saccades* respectively (Fig. 3 b). As shown in Fig. 3 c and d, the temporal structure of SS population responses differed between the two saccade directions consistently across adaptation and fatigue paradigms. For CS-on opposed saccades, PCs showed a sharp SS peak around saccade onset, whereas for CS-on aligned saccades, they exhibited a broad peak around saccade offset. To ensure comparable kinematics across both paradigms, only horizontal saccades from *baseline* phase of adaptation paradigm were included. Assuming that the CS-on aligned and CS-on opposed populations exert opposing influences on downstream targets, we estimated the collective impact of the two by calculating their difference (CS-on opposed – CS-on aligned, Fig. 3 e). This difference showed a robust biphasic profile with a sharp peak around saccade onset followed by a broad trough centering around 50 ms after saccade onset, a latency that approximately corresponds to the mean saccade duration. This peak-timing difference between CS-on opposed and aligned saccades was qualitatively consistent across both experimental paradigms, even when CF and CP saccades were analyzed separately (Suppl. Fig. 2 a,b), with comparable discharge patterns and peak/trough timings. A qualitatively similar biphasic SS population response was recently reported by Pi et al. (2024), providing independent support for the robustness of this profile. This reproducibility is noteworthy when considering the conspicuous cell-to-cell variability seen in PC SS discharge (De Zeeuw et al., 2023; Markanday et al., 2023; Sedaghat-Nejad et al., 2022; Tanaka et al., 2019), and is consistent with the notion that a PC-SS population signal rather than the individual SS-unit matters. It suggests that the difference profile may serve as a useful proxy for the collective impact of vermal PCs on downstream structures serving saccadic control. But what might such a signal encode?

If the SS population signal truly reflects an internal model’s estimate of actual movement kinematics, we reasoned that it should covary with the trial-by-trial variability observed in saccade kinematics, despite the same task demands within each condition. We therefore examined how SS firing rate scales with PV, a kinematic measure reflecting trial-by-trial variability. For each time bin, we regressed SS firing rate against the corresponding trial-by-trial PV values, yielding a time-resolved sensitivity profile expressed as the slope of the linear regression (in spikes/s per deg/s), where positive values indicate that higher PV was associated with higher SS firing rates and negative values indicate the opposite (Markanday et al. (2023), see Supplementary Methods for details). Fig. 3 f-h show mean PV-sensitivity profile aligned to saccade onset for CS-on opposed saccades, CS-on aligned saccades, and the difference between them, respectively. For CS-on opposed saccades, the PV sensitivity profile exhibited a biphasic modulation: a sharp peak around saccade onset, matching the timing of the peak of the SS firing rate, followed by a trough before saccade offset, indicating that the SS bursts reaches a higher peak and terminates earlier with saccades of higher PV (Fig. 3 f). In contrast, for CS-on aligned saccades, the PV sensitivity remained largely negative, showing a slight drop before saccade onset as SS firing rate started to rise, before remaining close to zero during the saccade. This indicates that higher PV was associated with lower SS firing rates before saccade onset, whereas during the movement, SS firing modulation was relatively little (Fig. 3 g). The difference profile (Fig. 3 h) shows an amplification of these profile features, characterized by a larger positive PV-sensitivity at saccade onset and a broader trough during and after the saccade, similar to the difference profile of the SS discharge in Fig. 3 e. This similarity, highlighted by the amplification of the key features in the difference profiles supports the idea that CS-on opposed and CS-on aligned SS units exert distinct influences on downstream targets in a coordinated, direction-dependent manner.

### Gain-changing adaptation involves direction-specific adjustments in SS population responses

Using this organizational framework for interpreting SS population responses, we then explored how gain-changing saccadic adaptation is reflected in the SS population signal. Since each PC was tested in only one adaptation direction, neither the outward nor inward adaptation group alone represents the full PC population needed to obtain a meaningful SS response profile for comparing the two forms of adaptation. Therefore, for visualization purposes, we reconstructed the net population SS activity after adaptation by adding the adaptation-induced change – computed separately for outward and inward adaptation as the mean difference between adaptation and baseline phases – to the total baseline population response including all recorded PCs. This approach preserves the full temporal structure of the population signal while illustrating how each form of adaptation modifies it. The adaptation-induced SS firing rate changes for each subpopulation are shown separately in Suppl. Fig. 3, and all statistical comparisons were performed on these subpopulation-specific changes rather than on the reconstructed population signal. Fig. 3 i and j show the reconstructed net population SS activities for CS-on opposed and CS-on aligned saccades, respectively, along with adaptation-induced changes expressed as effect size, separately for inward and outward adaptation sessions. The difference profiles are shown in Fig. 3 k. The results reveal clear differences that depend both on adaptation direction and saccade direction relative to the PC’s CS-on direction.

For CS-on opposed saccades, inward adaptation decreased the sharp peak of the firing rate around saccade onset compared to that of the baseline phase (mean reduction at saccade onset 5.5 ± 12.2 spikes/s; signed rank test; p=0.02), while outward adaptation produced a burst that lasted longer (mean increase at mean saccade offset, indicated by the vertical dashed line, 4.0 ± 12.5 spikes/s; signed rank test; p=0.02), with its peak height unchanged (mean increase at saccade onset 1.8 ± 8.6 spikes/s; signed rank test; p=0.28). Notably, this pattern of opposing changes between inward and outward adaptation mirrors the velocity profile changes observed behaviorally (Fig. 2 h), suggesting that CS-on opposed SS activity tracks the adaptation-induced changes in saccade kinematics. For CS-on aligned saccades, changes were smaller but in the opposite direction: outward adaptation slightly decreased SS firing rate at mean saccade offset (2.2 ± 16.46 spikes/s; signed rank test; p=0.43) and inward adaptation slightly increased it at saccade onset (1.7 ± 10.5 spikes/s; signed rand test; p=0.27). Although neither reached significance, these opposing changes amplified the CS-on opposed - CS-on aligned difference (Fig. 2 k) relative to CS-on opposed saccades alone (Fig. 2 i). Accordingly, the adaptation-induced changes were significantly amplified in the difference profile (Fig. 2 k): for outward adaptation (dashed green line), SS firing rate increased at mean saccade offset (6.2 ± 20.7 spikes/s; Welch’s t-test; p=0.023) and for inward adaptation (dashed purple line), it decreased at saccade onset (7.3 ± 16.1 spikes/s; Welch’s t-test; p=0.02), with the amplification confirmed by permutation testing (outward: p=0.009; inward: p=0.009).

To further examine whether these adaptation-induced SS changes are specific to the adapted direction, we computed the SS firing rate change for the diagonal non-adapted directions used to assess generalization. Since some sessions had very few trials for diagonal directions (Suppl. Fig. 1), reliable SS firing rates could not always be obtained for all directions. To address this, we exploited spatial continuity and fitted a cosine function over tested directions for each time bin, weighting by trial count as a reliability measure, separately for the baseline and generalization phase (see Supplementary Methods for detail). Heat maps of the adaptation-induced changes in effect sizes are shown in Suppl. Fig. 4 a-f, and changes at saccade onset and 50 ms after saccade onset are plotted as a function of target direction in Suppl. Fig. 4 g-i.

For CS-on opposed saccades, adaptation-induced changes gradually diminished with increasing angular deviation from the horizontal adapted direction, consistent with the behavioral generalization pattern. To quantify the directional tuning, we fitted a cosine function *d* = *acos* (*θ*−*ϕ*) + *c* to the effect sizes, where *a* reflects the gain and *ϕ* the phase of the tuning. For outward adaptation at mean saccade offset, this yielded *a*=0.91 and *ϕ*=12.6°, and for inward adaptation at saccade onset, *a*=-1.34 and *ϕ*=-12.5°. For CS-on aligned saccades, the corresponding fit yielded smaller and less consistent gains (*a*=-0.44, *ϕ*=- 9.3° for outward; *a*=2.06, *ϕ*=0.01 for inward), consistent with the weaker and non-significant changes observed for the horizontal direction. As with the horizontal direction, the CS-on opposed - CS-on aligned difference showed amplified tuning compared to CS-on opposed saccades alone, quantified by comparing the *in-phase component a*cos (*ϕ*): for outward adaptation this increase was minimal (0.10°; permutation test; 95% permutation null distribution: 0.90°, one-sided)., while for inward adaptation it was significant (1.67°; permutation test; 95% permutation null distribution: 1.47°, one-sided).

Finally, to assess whether similar SS population changes occur under fatigue, we compared SS firing rates between the first and last 50 trials of the *fatigue* paradigm (Suppl. Fig. 5). For CS-on opposed saccades, SS firing increased in the later part of the saccade (mean increase 2.2 ± 11.6 spikes/s at mean saccade offset; signed rank test; p=0.010), closely resembling the pattern observed during outward adaptation. In contrast, no significant change was observed for CS-on aligned saccades around movement. The parallel between fatigue-induced and outward adaptation-induced SS changes supports the interpretation that the cerebellum compensates fatigue-induced amplitude reduction through mechanisms that overlap with those underlying gain-increasing adaptation.

### CS around saccade onset showed directional sensitivity

Having demonstrated that SS population activity is modulated in a direction-dependent manner relative to the PC’s CS-on direction, we next asked whether a similar directional dependence is present in the CS response as well. We therefore examined CS population activity separately for horizontal CS-on opposed and CS-on aligned saccades. For the adaptation paradigm, horizontal saccades from all three phases were pooled to ensure sufficient CS counts. In contrast to SSs, the PC-CS population responses displayed highly similar task-related discharge patterns across the fatigue and the adaptation paradigms, and both rewarded CF and unrewarded CP saccades (Suppl. Fig. 2) with little difference between the two directions. The population CS firing rate started decreasing ∼100 ms before saccade onset, followed by two post-saccadic peaks, whose relative prominence differed between paradigms. In the case of the adaptation paradigm, two peaks were observed after saccade onset, whereas in the case of the fatigue paradigm, the first peak was much more prominent and the second peak was minimal. The two post-saccadic peaks have been reported to reflect distinct functions: the earlier peak, centered around 100–150 ms after saccade onset (i.e., ∼50-100 ms after saccade offset, given that a typical 15° saccade of duration of ∼50 ms), is associated with movement kinematics, putatively conveyed via proprioceptive feedback or efference copy (Markanday et al., 2021). Comparable CS responses to load-induced perturbation have also been reported during wrist movements (Gilbert and Thach, 1977). The later peak, around 200–250 ms after saccade onset (approximately 200 ms after saccade offset), reflects sensory error conveyed by visual feedback (Soetedjo et al., 2008). Consistent with this functional distinction, CS firing rate during the later post-saccadic error period showed greater modulation in the adaptation paradigm, where significant visual error was artificially introduced (Fig. 4 a), compared to the fatigue paradigm, where much smaller post-saccadic error arose from the natural variability of saccade endpoints (Fig. 4 b).

**Figure 4:**
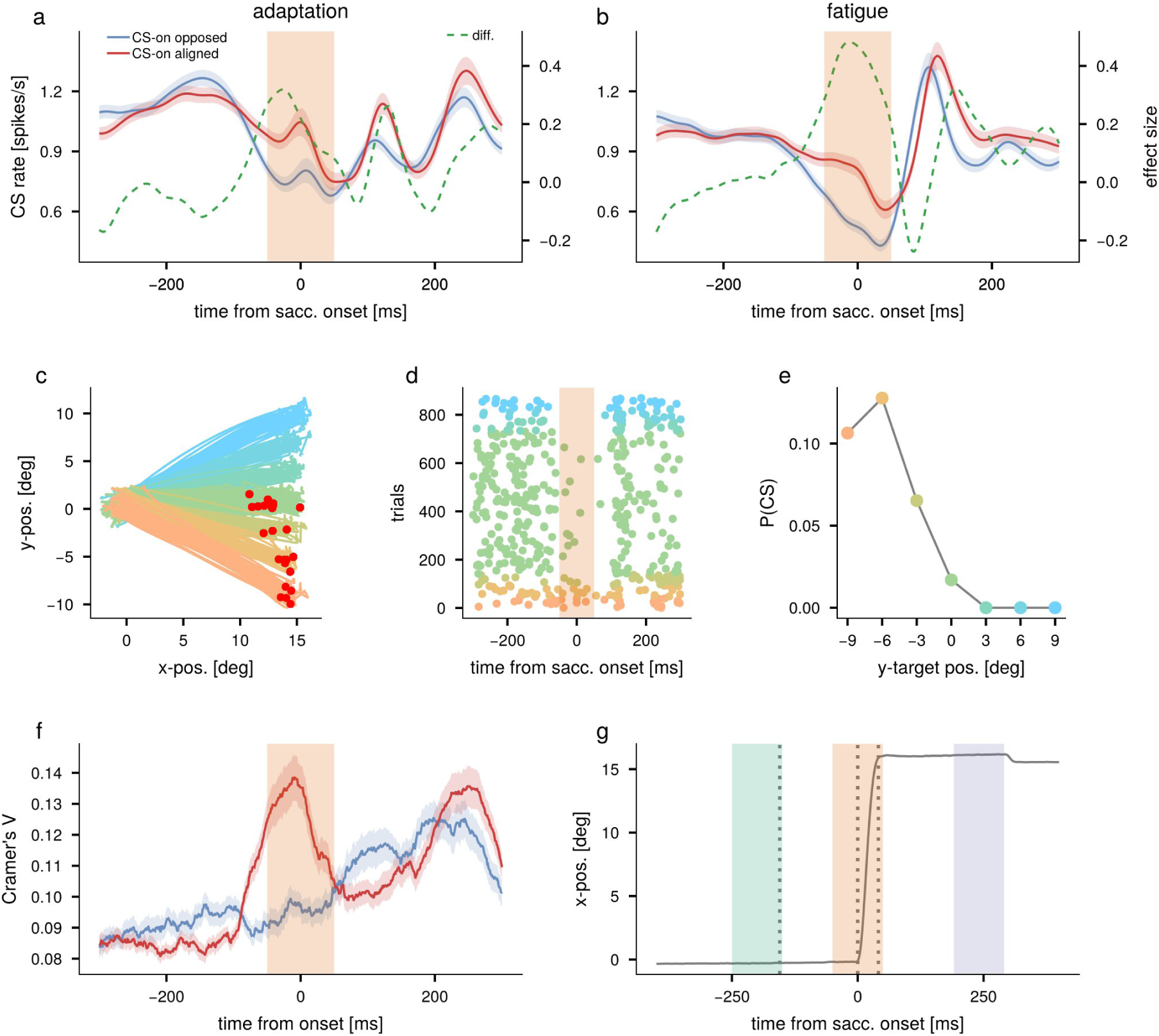
**a-b**: CS population responses for CS-on opposed and CS-on aligned saccades (a: adaptation paradigm, b: fatigue paradigm). The dashed green lines show the difference between the two directions as Cohen’s effect size. The orange shaded area indicates the peri-saccade onset period (-50 to 50 ms from onset). Shaded areas around traces indicate the standard error. **c-e**: CS activity during the peri-saccade onset period for an exemplary PC from an inward adaptation session. **c**: Eye position traces; red dots mark the saccade endpoint when a CS occurred during the peri-saccade onset period. Colors correspond to different target positions. **d**: Raster plot, with dot colors matching the same target positions in c. The shaded area indicates the peri-saccade onset period. **e**: CS firing probability during the peri-saccade onset period for each target position. **f**: Mean Cramér’s V computed between CS firing and target position for each PC subpopulation: CS-on opposed (blue) and CS-on aligned (red). **g**: Example trial illustrating three time windows of interest: *pre-saccadic error period* (-250 to -150 ms from saccade onset, green), *peri-saccade onset period* (-50 to 50 ms from saccade onset, orange), and post*-saccadic error period* (150 to 250 ms after saccade offset, purple). Dotted vertical lines indicate times of target onset, saccade onset, and saccade offset, respectively.

Focusing on time around saccade onset, the period in which the CS firing rate dropped, we found that the pre-saccadic decline in CS firing rate was attenuated for CS-on aligned compared to CS-on opposed saccades. The difference in CS firing rate between CS-on opposed and CS-on aligned saccades was largest around saccade onset. This tendency was consistent across both CF and CP saccades (Suppl. Fig. 2 c,d). To further investigate the functional role of CS firing around saccade onset, we examined the firing of individual CS units in relation to target location during the peri-saccade onset period (-50 to 50 ms relative to saccade onset; orange shaded area in Fig. 4 g). For this analysis, we focused on CF saccades from the adaptation paradigm, because the target appeared randomly at one of seven locations, whereas CP saccades were always directed to the center from seven different locations. We pooled trials across all three phases and all horizontal and diagonal target directions.

As exemplified in Fig. 4 c-e, the firing of some individual CS units during the peri-saccade onset period showed a clear dependence on target location, while others showed no such relationship. To examine which PCs showed directional selectivity around saccade onset, we quantified the strength of the directional preference of individual units in the peri-saccade onset period. To do this, we computed Cramer’s V, a measure of association between CS occurrence and target location, ranging from 0 (for no association) to 1 (for perfect selectivity). We computed Cramer’s V between occurrence of CS (present or absent) and 7 target locations for each time bin around saccade onset (see Supplementary Methods). The result, shown in Fig. 4 f, demonstrated that CS firing around saccade onset was associated with target locations only for CS-on aligned saccades, and moreover, that this significant association was confined to CF saccades and not seen in CP saccades (mean difference ± SD: CF, 0.030±0.022, p=0.0002; CP, 0.007±0.013, p=0.2036; Mann-Whitney U Test; Fig. 4 f and Suppl. Fig. 6). The interpretation of this difference between CP and CF saccades is complicated by the fact that the two saccade types differ in several respects. CP saccades were directed at a fixed midpoint and were effectively self-paced, as the lack of reward meant monkeys did not consistently wait for the go cue, nor were they required to execute them with high precision. Together, these factors gave CP saccades a clearly predictive and less demanding quality. CF saccades, by contrast, were directed at randomly varying target locations, required the monkey to wait for the go signal, and were rewarded only upon accurate fixation of the target.

### Effect of CSs during the peri-saccade onset period on ongoing saccade kinematics

To better understand the role of CSs that occur during the peri-saccade onset period, we divided saccades into two groups based on whether a CS was present (CS+) or absent (CS-; orange vs gray lines in Fig. 5), and compared the differences in their kinematic properties (dashed green lines in Fig. 5). For each kinematic parameter (PV, duration and amplitude), we fitted a linear mixed model with the kinematic parameter as the dependent variable, CS presence (binary) as a fixed effect, and session as a random effect, while accounting for saccade type (CF or CP), adaptation phase, and adaptation direction (see Supplementary Methods). For CS-on opposed saccades (Fig. 5 a, c and Suppl. Fig. 8 a, c), the presence of a CS during the peri-saccade onset period was associated with a reduction in PV (adaptation: −3.2 ± 1.2 deg/s, p=0.006; fatigue: −4.2 ± 1.5 deg/s, p=0.005). Despite this reduction in PV, the saccade amplitude was preserved (adaptation: 0.012 ± 0.02°, p=0.52; fatigue: -0.022 ± 0.016°, p=0.16), with duration significantly increased to compensate (adaptation: 0.38 ± 0.14 ms, p=0.006; fatigue: 0.29 ± 0.14 ms, p=0.043). The difference traces show how this compensation was achieved: the velocity reduction in the first half of the saccade was compensated by a comparable velocity increase in the second half, so that the position difference returned to near zero by the end of the movement. This CS-associated reduction in PV for CS-on opposed saccades is consistent with the finding of (Muller et al., 2023), who reported that CSs perturb ongoing eye movements toward the CS-on direction of the PCs, likely via the brief CS-induced pause in SS firing that transiently disinhibits DCN neurons. For CS-on aligned saccades (Fig. 5 b, d and Suppl. Fig. 8 b, d), eye position at saccade onset was significantly closer to the target on trials if a CS was present (adaptation: 0.046 ± 0.006°, p=7.4e-15; fatigue: 0.042 ± 0.008°, p=1.6e-7). Saccade amplitude was significantly reduced, preventing overshoot despite the closer starting position (adaptation: −0.076 ± 0.015°, p=3.2e-7; fatigue: −0.030 ± 0.014°, p=0.038), accomplished by shortening of duration (adaptation: −0.38 ± 0.10 ms, p=0.00012; fatigue: −0.54 ± 0.14 ms, p=0.00018) contrasting with the pattern observed for CS-on opposed saccades. The difference trace illustrates this: the positivie position difference before saccade onset was no longer present after the saccade because the velocity was reduced in the second half of the movement. For CS-on opposed saccades, we did not observe a comparable link between eye position at saccade onset and CS firing (adaptation: -0.012 ± 0.009°, p=0.15; fatigue: -0.002 ± 0.010°, p=0.85).

**Figure 5:**
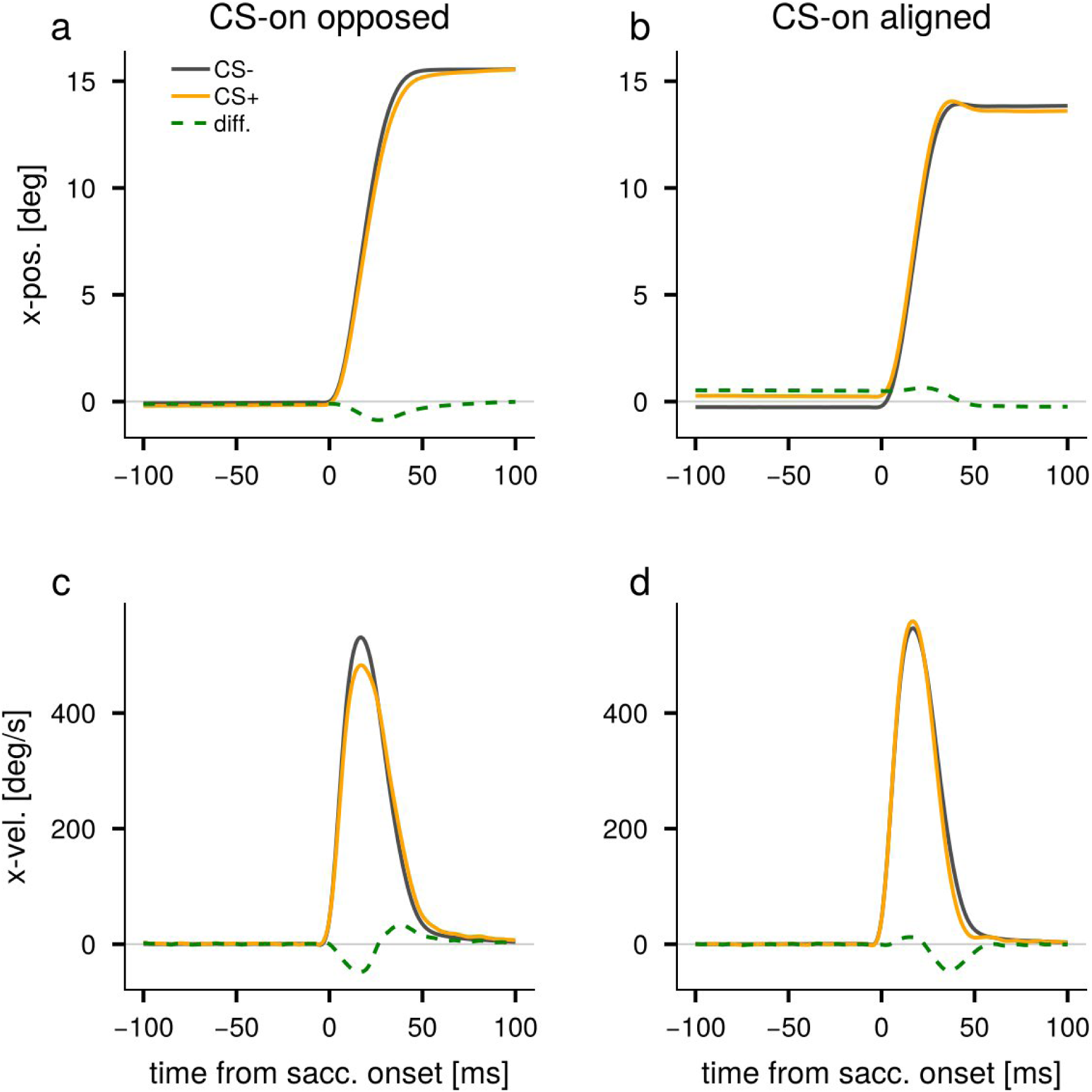
Effect of CS around saccade onset on saccade kinematics in the adaptation paradigm. **a-b**: Horizontal eye position when a CS occurred during the peri-saccade onset period (orange) and no CS occurred (gray), shown separately for CS-on opposed saccades (a) and CS-on aligned saccades (b). Dashed green lines show the difference. Note that the effect was amplified ten-fold for visualization. **c-d**: Same as a-b, but for horizontal eye velocity.

It is important to note that at saccade onset, the fixation point had already disappeared and the peripheral target was visible. The onset position-dependent CSs observed in CS- on aligned saccades are therefore fundamentally different from the visual error signals conventionally attributed to motor learning. For example, in a rightward saccade where the CS-on direction was also rightward, CSs fired more often when the eye was slightly deviated rightward from the central fixation point (Fig. 5 b) – indicating that CSs fired more for smaller retinal errors, the opposite of what would be expected for a conventional visual error signal. This suggests that the directional tuning of CS firing around saccade onset was opposite to that observed during the preceding fixation period. To test this, we analyzed CS modulation by fixation error (defined as eye position deviation from the central fixation point) during the pre-saccadic error period (-250 to -150 ms from saccade onset; green shaded area in Fig. 4 g) and the peri-saccade onset period. Indeed, the regression slopes quantifying CS modulation during the two periods showed opposite signs for CS-on aligned saccades (Fig. 6 d,f, Suppl. Fig. 9 d,f), confirming the tuning reversal. The regression slope during the peri-saccade onset period was significantly different from zero for CS-on aligned saccades (observed absolute slope: 0.11 spikes/s/deg; 95% permutation null distribution: 0.04 spikes/s/deg, one-sided) but not for CS-on opposed saccades. The finding that CS-on aligned CSs fire preferentially when the eye is closer to the target suggests that these CSs may play a role in motor planning rather than error correction –specifically, adjusting the upcoming saccade amplitude based on the eye’s starting position relative to the target, so that the movement is appropriately scaled.

**Figure 6:**
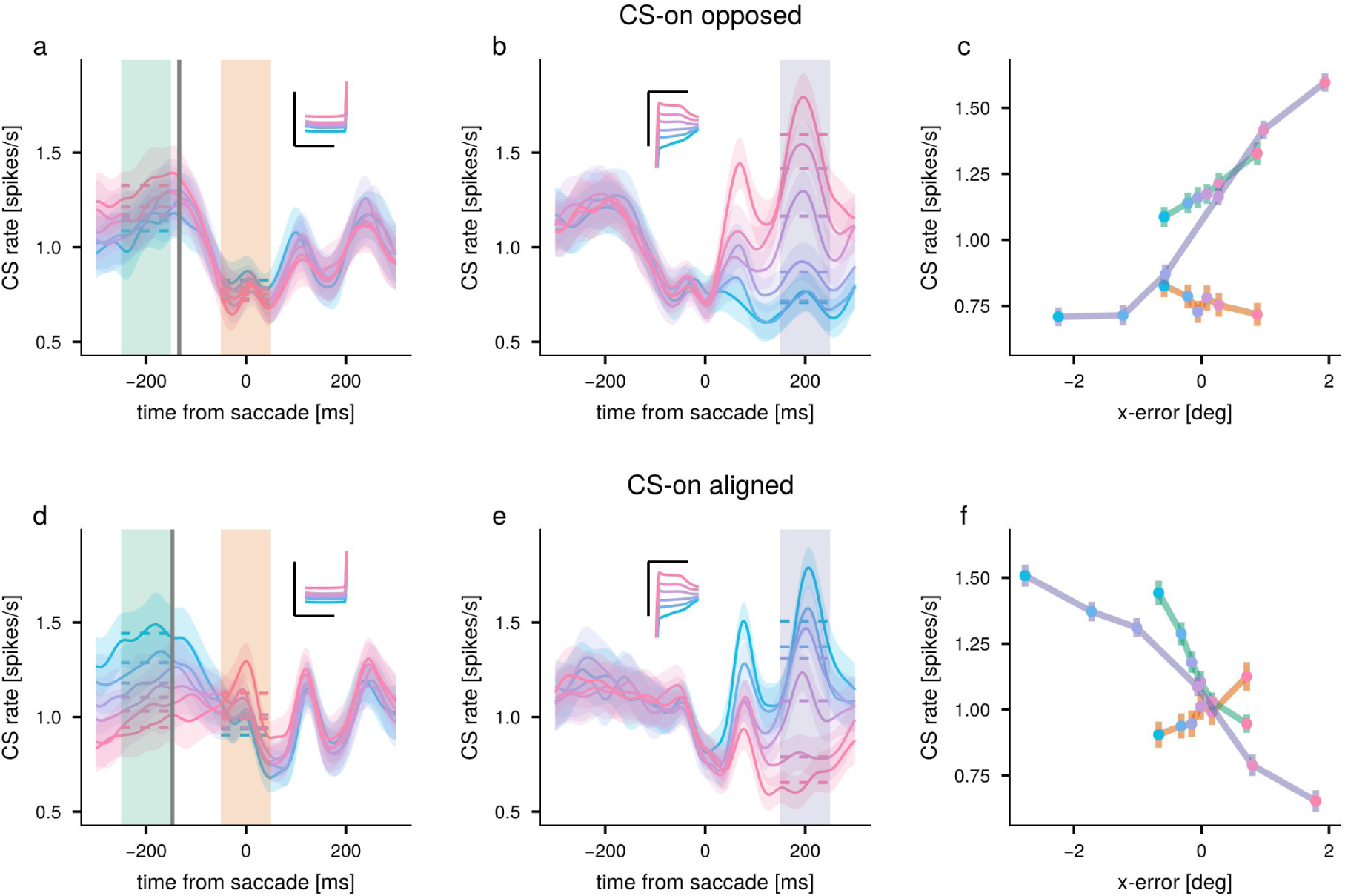
CS firing rate sorted by retinal error in the adaptation paradigm. **a**: Mean CS firing rate aligned to CS-on opposed saccade onset sorted by retinal error during the pre-saccadic error period (green shaded area). Dashed horizontal lines indicate the mean firing rate during the pre-saccadic error period and the perisaccade onset period (orange shaded area). The gray vertical line marks the mean time of the peripheral-target appearance. Inset: corresponding horizontal eye-position traces; scale bars indicate 5° and 300 ms. **b**: Same as a, but CS firing rates are aligned to CS-on opposed saccade offset. Dashed lines indicate the mean firing rate during the post-saccadic error period (purple shaded area). **c**: Mean CS firing rate during the three distinct periods as a function of retinal error. Line colors correspond to the shaded periods in a and b. d-f: Same as a-c, but for CS-on aligned saccades.

We then examined whether CS firing rates during the peri-saccade onset period were modulated by the adaptation state. For CS-on aligned saccades, CS firing rates did not change significantly between baseline and adaptation phases in either adaptation direction. In contrast, for CS-on opposed saccades, CS firing rate was significantly elevated during inward adaptation (signed rank test, p=0.02, Suppl. Fig. 7), a finding consistent with the earlier observation that peri-saccade onset CSs in CS-on opposed saccades reduce saccade velocity (Fig. 5 c), suggesting that elevated CS firing during inward adaptation may contribute to the kinematic signature of gain reduction – namely, the decrease in PV that characterizes inward adaptation (Fig. 2 h).

### CS error sensitivity is direction- and time-dependent

Having established that CS responses around saccade onset differ fundamentally from classical retinal error encoding, we next systematically characterized how CS error sensitivity varies over time around the saccade, extending the analysis to the post-saccadic error period (150 to 250 ms after saccade offset; purple shaded area in Fig. 4 g). In the post-saccadic error period, error was defined as eye position relative to the target, reflecting sensory prediction error arising from the intra-saccadic target displacement on adaptation trials, or from natural endpoint variability otherwise.

For CS-on opposed saccades, CS activity was only weakly modulated by fixation error in the pre-saccadic error period (green shaded area in Fig. 6 a for adaptation and Suppl. Fig. 9 a for fatigue), while it was strongly modulated in the post-saccadic error period (purple shaded area in Fig. 6 d for adaptation and Suppl. Fig. 9 d for fatigue), where CS firing reflected sensory prediction error. For CS-on aligned saccades, the pattern was inverted. CS activity was strongly modulated by fixation error in the pre-saccadic error pe-riod (Fig. 6 d for adaptation and Suppl. Fig. 9 d for fatigue), but this modulation was reduced in the post-saccadic error period (Fig. 6 d for adaptation and Suppl. Fig. 9 d for fatigue). Since targets were presented consistently on the same side throughout each session, the strong pre-saccadic modulation for CS-on aligned saccades suggests that anticipation of the upcoming target location toward the CS-on aligned direction enhances CS sensitivity to fixation error.

To quantify how error sensitivity changed from the pre- to post-saccadic error periods, we compared the regression slopes between the two periods (post-saccadic minus pre-saccadic error periods) separately for each direction. To control for the range of retinal errors for the adaptation paradigm, we limited this analysis to trials with post-saccadic error size < 1.5°, a threshold chosen to match the error range across conditions. For CS-on opposed saccades, error sensitivity increased from the pre- to the post-saccadic error period (adaptation: 0.35 spikes/s/deg; 95% permutation null distribution: [−0.10, 0.10]; fatigue: 0.16 spikes/s/deg; [−0.09, 0.08]; Fig. 6 c and Suppl. Fig. 9 f). In contrast, for CS- on aligned saccades, it decreased (adaptation: −0.13 spikes/s/deg; [−0.10, 0.10]; fatigue: −0.24 spikes/s/deg; [−0.08, 0.09]; Fig. 6 c and Suppl. Fig. 9 f). In all cases, the slope change fell outside the null distribution. This pattern was generally preserved across outward and inward adaptation sessions and for both CF and CP saccades (Suppl. Fig. 10, Suppl. Fig. 11).

Together, these results reveal a systematic and direction-dependent shift in CS error sensitivity across the three temporal windows. For CS-on opposed saccades, error sensitivity was lower in the pre-saccadic error period and increased in the post-saccadic error period, consistent with the conventional role of CS signals in driving motor learning through post-movement error correction. Around saccade onset, CS occurrence was associated with a reduction in PV and a compensatory prolongation of duration. For CS-on aligned saccades, error sensitivity was higher in the pre-saccadic error period, suggesting anticipatory modulation based on the upcoming target direction, and then declined in the post-saccadic error period. Around saccade onset, these saccades showed a tuning reversal, with CS firing reflecting the eye’s current state rather than retinal error and scaling the upcoming movement amplitude. This temporal reorganization suggests that the same PC population serves distinct computational roles across the stages of saccade planning, execution and error correction, depending on saccade direction.

### Peri-saccade onset CSs have similar effects on the SS population response despite different behavioral consequences

As described above, the direction-dependent kinematic effects of CSs during the peri-saccade onset period are likely mediated by their transient suppression of SS firing (∼20 ms pause following each CS; Markanday et al. (2020); Davie et al. (2008); Bloedel and Roberts (1971)), whose behavioral consequences differed qualitatively between CS-on aligned and CS-on opposed saccades. Given that the CS-on direction proved essential for interpreting SS population responses, we asked whether the different behavioral consequences of CSs would be reflected in corresponding different effects in the SS population activity, or whether CSs affect the SS population activity similarly in the two cases.

To evaluate the influence of CSs during the peri-saccade onset period on SS population activity, we compared the mean SS firing rates across three conditions: CS-on opposed saccades with a peri-saccade onset CS (CS+ opposed), CS-on aligned saccades with a peri-saccade onset CS (CS+ aligned), and saccades without a peri-saccade onset CS (CS-). In each condition, we calculated the SS rate difference between CS-on opposed and CS-on aligned saccades, and the conditions differed only in which trials were sampled. For CS+ opposed, only trials with a CS were included for CS-on opposed saccades and only trials without a CS for CS-on aligned saccades; for CS+ aligned, the arrangement was reversed; and for the control (CS-), only trials without a CS were included for both (Fig. 7 a). As shown in Fig. 7 b for adaptation and Suppl. Fig. 12 a for fatigue, the CS+ opposed condition produced a SS population response characterized by a double trough around saccade onset, while the CS+ aligned condition exhibited a prominent onset peak followed by a late minor trough.

**Figure 7:**
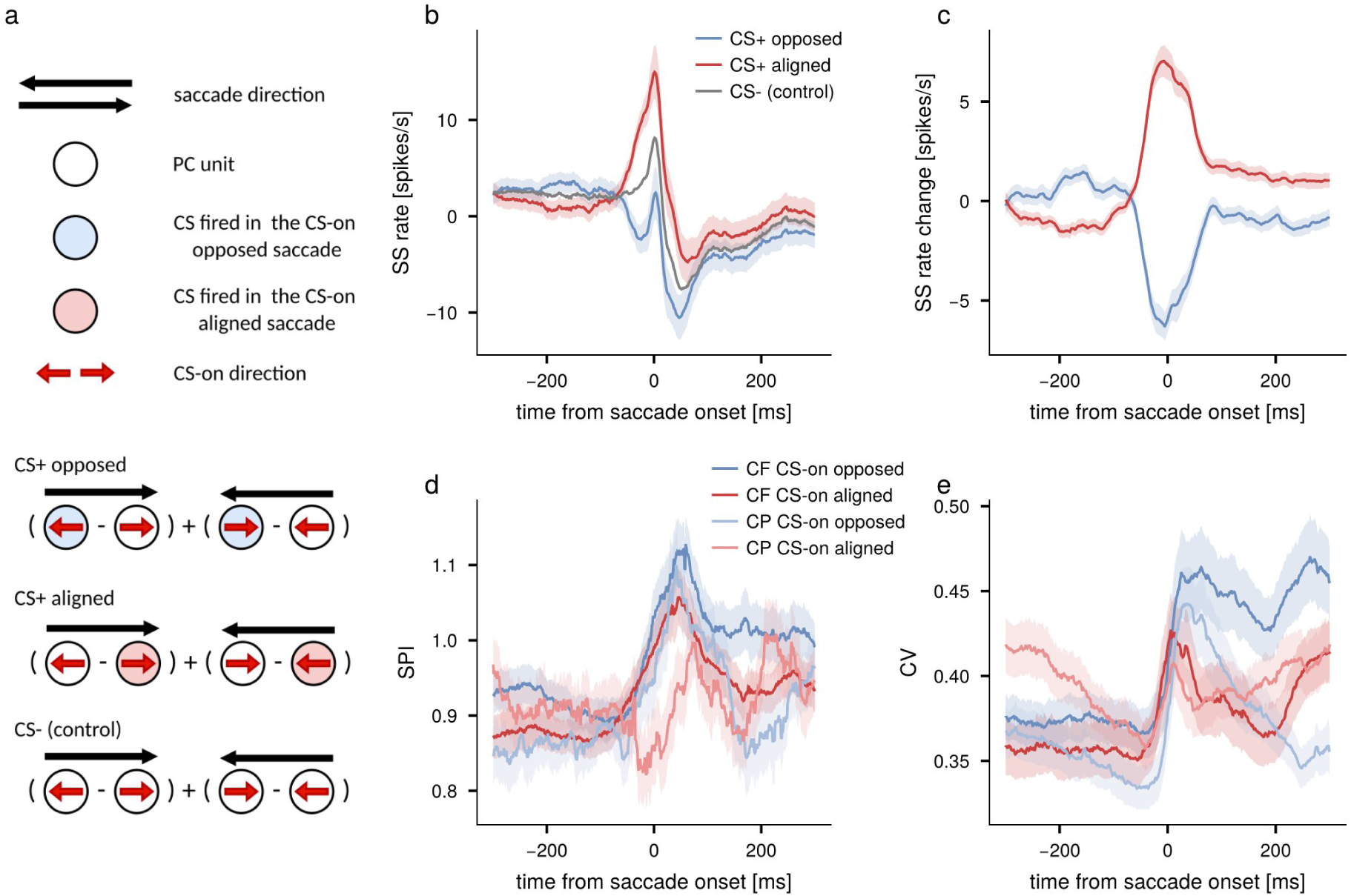
Effect of CS on net SS population responses in the fatigue paradigm. **a**: Schematic illustrating the three conditions used to assess the effect of CSs on SS firing. Each condition is the SS rate difference between CS-on opposed and CS-on aligned saccades, pooled over both saccade directions, with trials sampled according to the presence (color-filled circles) or absence (open circles) of a CS. **b**: Net SS population firing rate in the three conditions (blue: CS+ opposed, red: CS+ aligned, and gray: CS-). Shaded areas indicate the standard error. **c**: Changes in SS firing rate relative to the control conditions. **d**: Spike pause index (SPI) over time relative to saccade onset for CS-off and CS-on saccades, as well as for CF and CP saccades. Shaded areas indicate the standard error. **e**: Same as d, but for coefficient of variation (CV).

Despite these qualitatively distinct SS modulation patterns, the CS-induced change in the net SS population was comparable in magnitude for the two conditions: CSs in the CS+ aligned condition increased the net population SS firing rate relative to CS-control condition, while those in the CS+ opposed condition decreased it by a comparable amount (Fig. 7 c for adaptation and Suppl. Fig. 12 b for fatigue). The opposite signs are a consequence of calculating the population signal by subtracting the CS- control condition from CS+ opposed and CS+ aligned condition. This suggests that the same CS-induced suppression of SS firing lowers the signal when it occurs in one population and raises it when it occurs in the other. The duration of this CS-induced modulation closely overlapped with the biphasic SS population response described earlier, (Fig. 3 e), suggesting that peri-saccade onset CSs contribute to shaping the overall SS population signal. This pattern was consistent across both experimental paradigms. Together, these results suggest that peri-saccade onset CSs suppress SS firing to a similar extent in the two subpopulations. In contrast, the behavioral consequences of these CSs differed with saccade direction relative to the PC’s CS-on direction.

### Duration of CS-induced SS pause varies across the peri-saccadic period

While CSs induce a brief pause in SS firing, the duration of this pause may depend on when the CS occurs relative to the saccade. We therefore asked whether the duration of the CS-induced SS pause varied systematically across the peri-saccadic period. To quantify the duration of the CS-induced SS pause, we introduced a spike pause index (SPI; see Supplementary Methods), defined for the *i*^th^ CS as:

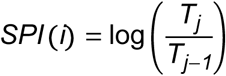

where *T_j_* is the *i*^th^ interspike interval (ISI) of SS that contains the *i*^th^ CS. The SPI quantifies the extent of the pause by comparing the post-CS ISI (*T_j_*) with the pre-CS ISI (*T_j_*_−_*_1_*). A large SPI indicates a prolonged pause, while an SPI of zero indicates no change. SPI traces were smoothed using a moving window with a 50 ms half-width.

The SPI increased from saccade onset toward saccade offset, peaking around offset (∼50ms), consistently across CS-on aligned and CS-on opposed saccades, for both CF and CP saccades, and in both paradigms (Fig. 7 d for adaptation and Suppl. Fig. 12 c for fatigue). While the baseline SPI before saccade onset varied modestly across 4 conditions (CS-on aligned/opposed, CF/CP saccades), the increase from its pre-saccadic baseline to its peak around saccade offset did not differ among them (linear mixed model; CS-on aligned/opposed: p=0.84 for adaptation, p=0.17 for fatigue; CF/CP: p=0.76 and p=0.89: interaction: p=0.088 and p=0.050). The consistency of this temporal pattern is notable given considerable cell-to-cell variability in SS activity (De Zeeuw et al., 2023; Markanday et al., 2023; Sedaghat-Nejad et al., 2022; Tanaka et al., 2019). Following saccade offset, the diverging SPI trajectories in the adaptation paradigm – compared to the steady decrease seen in the fatigue paradigm – likely reflect the influence of corrective saccades.

To identify the source of this SPI modulation, we considered two candidate explanations. First, longer pauses might simply reflect longer CS events (i.e., a change in the climbing fiber input itself), which is why we tested whether SPI was related to CS duration. Second, as we will elaborate in the Discussion, a longer pause could reflect a change in the synaptic input to PCs, namely the MF-PF pathway and the MLIs it drives: stronger, noisier excitatory and inhibitory drive increases trial-to-trial variability in SS firing and may extend the suppression of SS firing after a CS. We therefore used the coefficient of variation (CV, SD/mean) of SS firing as an index of this variability. CS duration was not significantly correlated with SPI, with correlation coefficient near zero (r = 0.0056 ± 0.0054 SEM for adaptation and r = -0.0014 ± 0.0062 SEM for fatigue). In contrast, the CV of SS firing rate was strongly correlated with SPI (r = 0.84, 95% bootstrap null distribution: [0.47,0.95] for adaptation and r = 0.88, 95% bootstrap null distribution: [0.61,0.97] for the fatigue). The CV increased shortly before saccade onset and peaked around saccade offset, mirroring the SPI profile, and this pattern was consistent across all PC subpopulations and saccade directions (Fig. 7 e for adaptation and Suppl. Fig. 12 d for fatigue). Together, these results indicate that the duration of the CS-induced SS pause is dynamically modulated around saccadic movement and that this modulation is more closely associated with the CV of SS firing than with CS duration itself. The close parallel between the SPI modulation and the peri-saccadic rise in CV suggests a common underlying mechanism, which we consider in the Discussion.

## Discussion

Our study confirmed and extended previous findings that cerebellar PCs in the OMV are spatially organized according to their preference for the direction of post-saccadic retinal error and that this organization provides a viable framework for interpreting SS population activity and its behavioral consequences (Fakharian et al., 2025; Herzfeld et al., 2015; Pi et al., 2024). Our data indicate that CS-on aligned and CS-on opposed subpopulations are not merely mirror images of one another but exhibit features that suggest functionally distinct roles in shaping the saccadic motor command. The CS-on opposed subpopulation of PCs exhibits a precisely timed sharp peak of SS activity at saccade onset that scales with PV, while the CS-on aligned subpopulation exhibits a broader SS modulation around saccade offset. Beyond their distinct SS discharge profiles, the two PC subpopulations also differed in their CS error sensitivity: error encoding was stronger for the CS- on aligned subpopulation during the pre-saccadic error period, while CS-on opposed subpopulation showed stronger encoding during the post-saccadic error period. Furthermore, peri-saccade onset CS firing had direction-dependent kinematic consequences, and the duration of the CS-induced SS pause was dynamically modulated across the peri-saccadic period, correlating with the CV of SS firing rather than CS duration. Together, these findings provide direct electrophysiological evidence that the CS-on directional preferences of individual PCs are a critical determinant of their functional contribution to saccadic control. In the following, we discuss how these findings inform our understanding of the distinct computational roles of forward and inverse models in saccadic control, and why combining both control strategies within the same PC population may confer specific functional advantages.

### Movement direction-dependent functions of PCs

Our results reveal that PCs play functionally distinct roles depending on whether the saccade direction is aligned or opposed to their CS-on on direction, and that this functional distinction maps onto the spatial organization of the OMV based on the CS-on directions.

For CS-on aligned saccades, several lines of evidence suggest that these PCs contribute primarily to movement planning in a feedforward manner. First, CSs were more sensitive to retinal error during the pre-saccadic error period than during the post-saccadic error period (Fig. 6, Suppl. Fig. 9), indicating that this subpopulation monitors the state of the eye before the movement rather than correcting it afterward. Second, peri-saccade onset CSs in this subpopulation encoded both target location (Fig. 4 c-f) and eye position at saccade onset (Fig. 5 b, Suppl. Fig. 8 b), consistent with a role in specifying the initial conditions for the upcoming movement. Third, SS firing was not modulated by PV during the saccade itself, but showed PV sensitivity shortly before saccade onset (Fig. 3 g), suggesting that the motor command is pre-programmed rather than updated online. A final argument builds on the well-established inhibitory control of cFN neurons by vermal PCs (Fig. 8 a). The broad SS burst centered around saccade offset in CS-on aligned PCs suppresses the contraversive cFN during the later phase, allowing it to fire predominantly in the early phase and thereby drive agonist saccadic burst neuron (SBN) activity during acceleration (Fig. 8 b) – the full circuit logic is elaborated below. Together, these properties are consistent with an inverse model operating in a feedforward pathway: a controller that computes and issues the motor command based on the planned movement before execution begins.

**Figure 8:**
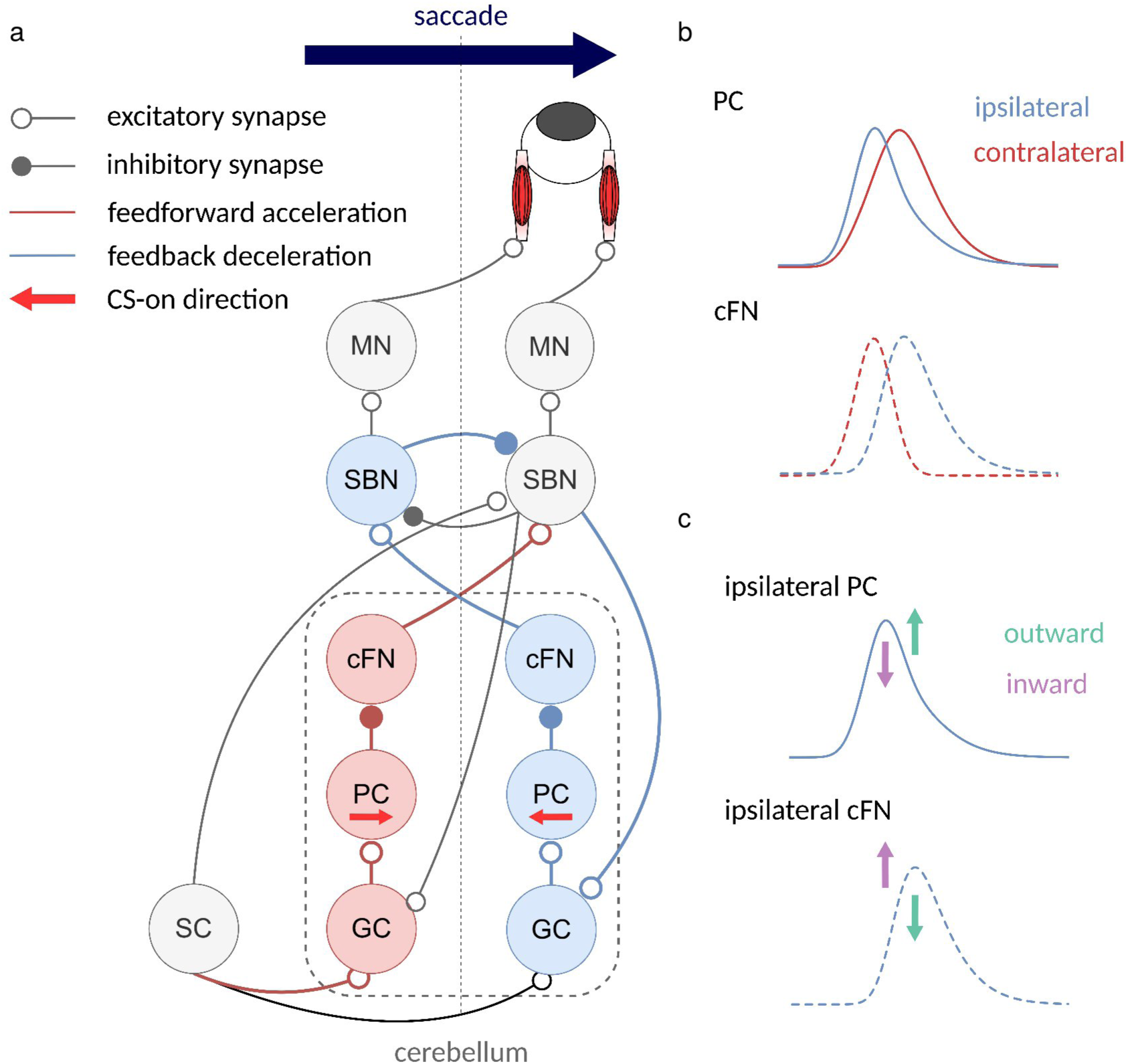
**a**: Simplified schematic of a proposed circuit. PCs ipsiversive to agonist saccadic burst neurons (SBN) receive input from them and project via the ipsiversive cFN to the antagonist SBN, forming a negative feedback loop that contributes to the saccade deceleration (blue pathway). Conversely, PCs contraversive to the movement direction disinhibit the agonist SBN via the contraversive cFN, accelerating the eyes based on target displacement information available prior to movement onset (red pathway). MN: motoneuron; SC: superior colliculus. **b**: Schematic illustration of how the early and late SS burst of PCs may shape their down-stream cFN activity. An early burst in ipsiversive PCs suppresses early firing in the ipsiversive cFN, allowing them to fire only later, whereas a late burst in contraversive PCs suppresses the later cFN firing, allowing them to fire only during the acceleration phase. **c**: Possible mechanism underlying gain-changing adaptation. During outward adaptation, the later phase of the SS burst in ipsiversive PCs increases, reducing later firing in ipsiversive cFN; this delays saccade termination, producing larger amplitudes and longer durations. During inward adaptation, the early SS burst in ipsiversive PCs is reduced, leading to enhanced early burst in the ipsiversive cFN, which decreases PV and reduces amplitude.

For CS-on opposed saccades, by contrast, the evidence points to a complementary role in online movement control. Here the CSs were more sensitive to retinal error during the post-saccadic error period than to retinal error during the pre-saccadic error period. This indicates that this subpopulation prioritizes post-movement error signals that reflect the accuracy of the movement just completed – the type of error signal needed to update the sensory predictions of a forward model. Supporting this, SS firing showed robust PV sensitivity during the saccade, with higher PVs producing a more pronounced but temporally narrower SS burst (Fig. 3 f), indicating that this subpopulation monitors ongoing movement kinematics in real time. Furthermore, peri-saccade onset CSs in this subpopulation were associated with reduced PV, consistent with the movement being transiently pulled toward the CS-on direction (Muller et al., 2023), while duration was prolonged to maintain amplitude, consistent with an online corrective signal. The early sharp SS burst may suppress ipsiversive cFN firing during the early phase, allowing it to fire later to dynamically adjust deceleration (Fig. 8 a, b) – consistent with the circuit organization discussed below. Additionally, SS firing in this subpopulation was changed after adaptation; outward adaptation prolonged the burst, while inward adaptation reduced its peak, reflecting the updated kinematic features of the adapted saccade. This is what would be expected of a forward model that is updated by post-movement sensory prediction error signals to produce more accurate state estimates for future movements (Fig. 8 c). Together, these properties are consistent with a forward model operating within a feedback loop: a controller that estimates the current state of the movement in real time and signals deviations from the intended trajectory to downstream structures for online correction.

Since CS-on aligned PCs are predominantly located in the contraversive OMV and CS-on opposed PCS in the ipsiversive OMV, these functional distinctions map directly onto anatomically distinct cerebellar domains, consistent with the optogenetic perturbation results of (Soetedjo and Horwitz, 2024), who demonstrated differential effects of ipsiversive and contraversive cortex stimulation during saccades. The circuit logic underlying this organization is indirect and not immediately intuitive, owing to the inhibitory nature of PC output and the contralateral projection of cFN neurons (Fujita et al., 2020), which means that PC activity is sign-inverted twice before reaching the motor output. As a result, the early sharp SS burst in ipsiversive PCs suppresses ipsiversive cFN firing to modulate deceleration, while the later broad burst in contraversive PCs suppresses contraversive cFN activity to facilitate early acceleration by disinhibiting downstream premotor neurons – giving rise to a temporally precise, opponent modulation of cFN output that jointly shapes the acceleration-deceleration profile of the movement (Fig. 8 a).

### Effect of adaptation and fatigue on SS

A notable asymmetry emerged between inward and outward adaptation in both behavior and SS firing. During inward adaptation, the decrease in saccade amplitude was accompanied by a reduction in PV, and the SS peak firing rate decreased accordingly. In contrast, during outward adaptation, amplitude increases were achieved primarily through prolonged saccade duration rather than increased PV and SS firing increased during the later phase of the movement while the PV remained largely unchanged.

This asymmetry may reflect the contribution of non-motor fatigue to saccadic adaptation as a major mechanism underlying inward adaptation. A previous study showed that patients with pathology involving the cerebellar vermis can adapt their saccades inward but not outward following repetitive intra-saccadic shifts (Golla et al. (2008)), suggesting that fatigue, a term that comprises the inevitable loss of vigor due to muscular and/ or cognitive exhaustion is exploited to realize amplitude reduction demanded by inward adaptation paradigms. Of course normal saccades are also subject to fatigue and the consequential drop of saccade velocity. In healthy subjects, however, amplitude is maintained despite fatigue-induced PV reduction by a compensatory extension of duration. This compensation appears to be absent in the patient group, whose saccade amplitude progressively decreased during longer saccade sequences. Outward adaptation may exploit the same duration-extending mechanism, increasing amplitude beyond what is needed to merely compensate for fatigue-induced velocity loss. Consistent with this interpretation, the SS firing changes induced by fatigue in our data closely resembled those observed during outward adaptation, with increases most prominent in the later portion of the saccade in the CS-on opposed subpopulation (Suppl. Fig. 5). Together, these findings suggest that the cerebellum continuously compensates for fatigue induced amplitude reduction through mechanisms that overlap with those underlying outward gain-changing adaptation.

### Role of CSs around saccade onset

Having characterized the distinct functional roles of CS-on aligned and CS-on opposed subpopulations of PCs in terms of their SS discharge and error sensitivity, we next consider the specific contribution of peri-saccade onset CS firing to movement control.

For CS-on opposed saccades, peri-saccade onset CSs reduced PV while saccade duration was prolonged to preserve endpoint accuracy. This kinematic pattern can be explained by the circuit consequences of CS firing: the transient CS-induced pause in ipsiversive SS activity disinhibits ipsiversive cFN neurons, increasing their contralateral excitatory output to antagonist SBNs, thereby enhancing the braking drive opposing the ongoing movement. Consistent with a role in online movement control, CS firing rate during the peri-saccade onset period was increased during inward adaptation – in line with the behavioral reduction in PV – while remaining unchanged during outward adaptation, consistent with the forward model interpretation discussed above.

For CS-on aligned saccades, peri-saccade onset CS firing was dependent on eye position at saccade onset, occurring more frequently when the eyes were already displaced closer to the target. Because this displacement reduces the required saccade amplitude, the observed amplitude reduction associated with peri-saccade onset CS firing can be understood as an appropriate fine adjustment of movement size based on the initial eye position, preventing overshoot despite the closer starting position. However, the circuit mechanism underlying this adjustment is not immediately apparent. The CS-induced SS pause in contraversive PCs would disinhibit contraversive cFN neurons, which would be expected to enhance agonistic drive and increase rather than decrease amplitude. Furthermore, the regression slope between CS firing and retinal error during the peri-saccade onset was reversed relative to both the pre-saccadic and post-saccadic error periods, suggesting that these CSs do not simply encode classical visual error. The mechanistic link between peri-saccade onset CS firing in CS-on aligned saccades and the resulting amplitude reduction therefore remains an important open question and may involve interactions between the cerebellar output and upstream premotor circuits that are not captured by the current analysis.

### CS-induced SS pause is dynamically modulated around saccadic movements

Finally, beyond their directional specificity, CS-induced SS pauses were themselves dynamically modulated across the peri-saccadic period. As shown in Fig. 7 d, the duration of the CS-induced SS pause increased shortly before saccade onset and peaked around saccade offset, a pattern that was consistent across all PC subpopulations regardless of individual SS discharge patterns or saccades directed aligned with or opposed to the CS- on direction.

The close parallel between the modulation of pause duration captured by the SPI measure and the peri-saccadic rise in CV in SS firing suggests a common underlying mechanism. A concurrent increase in both excitatory and inhibitory synaptic inputs with independent noise to PCs likely reflecting the increased parallel fiber and MLI drive associated with active movement – a condition known to increase firing variability (Shadlen and Newsome, 1998) – would naturally produce both a higher CV and a prolonged post-CS pause, as the inhibitory input would extend the suppression of SS firing following each CS. This interpretation is further supported by the demonstration of (Mathews et al., 2012) that the duration of the post-CS SS pause is modulated by input from inhibitory MLIs. Consistent with this, (Sedaghat-Nejad et al., 2022) reported increased SS synchronization among OMV PCs during the deceleration phase of saccades – a finding that coincides with the timing of our SPI peak around saccade offset – and similarly proposed that MLI input may regulate the degree of synchronization. Together, these findings suggest that the dynamic modulation of CS-induced SS pauses during saccades reflects a peri-saccadic increase in inhibitory drive to PCs from MLIs, which extends the suppression of SS firing following each CS and may thereby amplify the functional impact of the transient disinhibition of downstream cFN neurons.

## Conclusion

The low discharge rate of CSs has long suggested a comparatively simple functional role – a sparse error signal driving synaptic plasticity at PF-PC synapses. However, growing evidence that CS duration, waveform morphology, and precise timing carry information beyond a simple binary error signal has prompted a reconsideration of this view. The present study adds a further dimension: by considering the geometric relationship between CS-on direction and movement direction, we show that individual PCs can multiplex a variety of information streams across time – monitoring pre-movement eye state, signaling online kinematic perturbations, and encoding post-movement sensory prediction errors – with their functional contribution depending critically on the direction of the ongoing saccade relative to their CS-on direction.

Why might the cerebellum combine forward and inverse model computations within the same PC population? Saccades face an inherent speed-accuracy trade-off: they must be executed rapidly – within tens of milliseconds – yet land with high endpoint precision. We propose that the two types of internal models serve complementary roles in resolving this trade-off: the inverse model drives the high initial acceleration needed for a fast movement, while the forward model monitors the ongoing trajectory and secures endpoint precision by anticipating and correcting deviations in real time. Implementing both within a single, spatially organized population allows this coordination to occur within minimal communication delay – a critical advantage given the brevity of the saccadic movement itself.

Importantly, we do not propose that forward and inverse model computations are strictly modular or that the OMV is divided into two clearly segregated functional zones. Indeed, the differences in CS error sensitivity between the pre- and post-saccadic periods were graded rather than absolute – both subpopulations retained some sensitivity to both error periods, differing in the degree rather than the type of error encoding. This graded sensitivity, combined with the fact that PC SS firing patterns exist along a continuum of bursts, pauses, and their combinations, and that the OMV lacks a sharp anatomical boundary between its left and right halves, suggests that the functional distinction between forward and inverse model computations reflects a continuous gradation across the PC population rather than a strict categorical division (Markanday et al., 2023). As saccades deviate from the horizontal axis into diagonal or vertical directions, the contribution of each PC to forward and inverse model computations likely shifts gradually, reflecting the continuous nature of the underlying population code. This continuum of functionality, embedded within a compact and spatially organized circuit, may represent a general principle by which the cerebellum achieves flexible, context-dependent motor control – one that extends beyond saccades to other movement systems where speed and precision must be simultaneously optimized.

## Supplementary methods

### Animals, preparation, surgical procedures, and recording methods

Data were recorded from three adult male rhesus macaques (Macaca mulatta): Monkey K (10 years), Monkey E (8 years), and Monkey I (10 years). The fatigue paradigm dataset, involving Monkeys K and E, was previously reported in (Markanday et al., 2023, 2021). For the adaptation paradigm, Monkeys K, E, and I were utilized. All experimental, training, and surgical protocols were conducted in accordance with German and European law and the National Institutes of Health’s Guide for the Care and Use of Laboratory Animals. All procedures were approved and supervised by the local authority (Regierungspräsidium Tübingen) under veterinary licenses N7/18 and N4/14. The surgical and training procedures for the adaptation paradigm were identical to those established for the fatigue paradigm (see Markanday et al. 2021 for comprehensive details). Briefly, animals were habituated to a primate chair before undergoing a multi-stage surgical process. Titanium foundations were first secured to the skull and allowed to osseointegrate for 3–4 months. Subsequently, a head post for stable immobilization and magnetic scleral search coils for high-resolution eye-tracking were implanted. Following behavioral training, a titanium recording chamber was positioned over the cerebellar midline, tilted 30° posteriorly. This orientation, planned and confirmed via pre- and post-surgical MRI, provided electrode access to the OMV (lobules VIc/VIIa). All surgeries were performed under aseptic conditions using general anesthesia while vital physiological parameters (heart rate, blood pressure, pO₂, pCO₂, and body temperature) were continuously monitored. Post-operative analgesia was provided via buprenorphine, and recovery was assessed through regular ethograms. Animals were allowed to recover fully before the commencement of experimental sessions.

### CS detection and post-processing

CS detection and post-processing procedures were identical to those described in (Markanday et al., 2023, 2021). Briefly, CSs were detected using a deep neural network classifier trained to identify complex spike forms, which has been shown to outperform conventional template matching algorithms in terms of sensitivity and accuracy (Markanday et al., 2020). In addition to detecting CS occurrence, the network also provided estimates of individual CS duration, capturing the temporal extent of each CS waveform. Post-processing steps, including false positive exclusion, followed the same criteria as in (Markanday et al., 2023, 2021).

### Defining CS-on direction

To determine each PC’s CS-on direction, we analyzed CS firing rate during the post-saccadic error period (50 to 250 ms after saccade offset), independent of the saccade direction. To remove endpoint bias, we subtracted the median retinal error (the difference between the saccade endpoint and the target) separately for CF and CP saccades. The corrected errors were then divided into two bins of the equal size (leftward and rightward errors) and these bins were used to compute the CS probabilities associated with each error direction. The side with the higher CS probability during this period was defined as the CS-on direction of that PC.

### Firing rate estimation

A common approach to estimating instantaneous firing rates is to smooth spike trains with a Gaussian kernel. However, selecting an appropriate kernel width requires balancing temporal resolution against smoothness: a narrow kernel preserves rapid changes of the signal but yields a noisy estimate, whereas a wide kernel produces a smoother signal at the cost of temporal precision. To avoid this trade-off, we estimated SS firing rates using a maximum a posteriori (MAP) estimator under linear Gaussian assumption, which preserves temporal resolution while reducing noise.

The key idea is that a single trial provides limited information about the underlying firing rate, but the mean firing rate across trials can serve as a prior, because our tasks involve the highly repetitive movements. With a sufficient number of trials, this mean rate becomes a smooth estimate even without explicit kernel smoothing. The MAP estimator combines this prior with the single-trial spike train, yielding a trial-specific firing-rate estimate that is both smooth and temporally precise. Conceptually, this approach is equivalent to a Kalman filter: the trial-averaged firing rate acts as a prior, and the single-trial spike train provides the observation, allowing the estimator to combine both sources of information optimally.

For this smoothing procedure, we required an observation matrix *x* (with *n* observations by *l* time bins), a transition vector *u* (length *l*−*1*), an observation-uncertainty vector *v_x_* (length *l*), and a transition-uncertainty vector *v_u_* (length *l*−*1*). We used inverse interspike interval (IISI), aligned to saccades, for *x*. The transition vector *u* was defined as the gradient of the trial-averaged *x*, while the observation uncertainty *v_x_* was taken as the variance of *x*. The transition uncertainty *v_u_* was defined as the variance of *u*. The gradient *u* at time *i* was computed as:

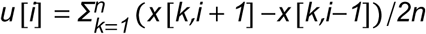

The estimation procedure consists of two steps; forward pass (filtering) and backward pass (smoothing). The forward pass incorporates observations from time 1 through *i*, whereas the backward pass carries information from future observations. Together, these two passes ensure that the estimate at time *i* reflects the entire observation sequence.

For the forward pass, let *y_f_* [*k*,*i*] denote the filtered state at time *i* of the *k*-th trial (*i* > *1*). We initialized

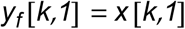

The update equation is

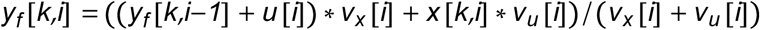

This recursion is applied sequentially until *i* = *l*. For the backward passs, let *y_s_* [*k*,*i*] denote the smoothed state at time *i* of the *k*-th trial (*i* < *l*). We initialized

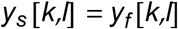

The update equation is

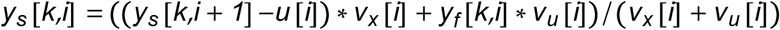

This recursion is applied sequentially until *i* = *1*. This MAP smoothing method also allowed to derive a smoother estimate PV sensitivity over time compared with conventional kernel smoothing. Suppl. Fig. 13 illustrates results from two exemplary cells. To evaluate the performance, we compared this approach with four other smoothing methods: raw IISI, IISI smoothed with a Gaussian kernel (SD = 5 ms), and spike trains smoothed with Gaussian kernels (SD = 5 ms and 10 ms). As shown in Suppl. Fig. 13, MAP estimator effectively smoothed both single-trial firing rates and PV sensitivity. In contrast, although wider Gaussian kernels smoothed the firing rate, they also distorted the sharp peak in the mean firing rate. The main limitation of this method is the underlying linear-Gaussian assumption. The distribution of IISI was treated as approximately linear and Gaussian, but when the firing rates are low, the distribution becomes increasingly skewed. However, SSs exhibit sufficiently high baseline firing rates that this approximation remains reasonable for our data. In contrast, CSs fire at extremely low rates, making this MAP estimation method inappropriate for these events. Therefore, we instead applied a conventional Gaussian kernel with a 15 ms SD.

For all subsequent population-level analyses, SS firing rates were baseline-corrected by subtracting the mean firing rate during the period from -250 to -150 ms from saccade onset, computed separately for each paradigm phase. This correction was applied to individual PC units prior to computing subpopulation means for CS-on opposed and CS-on aligned saccades, as well as for quantifying adaptation-induced changes.

### Computing PV sensitivity

To compute SS sensitivity to PV, the slope of a linear regression between SS firing rate and PV was computed for every 1 ms time bin. For the fatigue paradigm, this regression was computed directly. For the adaptation paradigm, because the baseline phase alone contained an insufficient number of trials for reliable slope estimation, trials were pooled across all three paradigm phases. A linear model was used to account for these phase-specific differences while estimating the overall PV sensitivity:

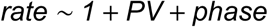

where *PV* captures the population-level PV sensitivity and *phase* accounts for differences in baseline firing rate and kinematics across the three adaptation phases. The PV sensitivity at each time bin was extracted from the coefficient of PV.

### Fitting cosine function

Since some sessions had insufficient trials in diagonal directions to reliably estimate SS firing rates (Suppl. Fig. 1), we exploited the spatial continuity of saccade directions to obtain stable estimates across all directions. For each time bin *t*, a cosine function was fitted to the observed firing rates across the seven target directions using weighted least squares, with the number of trials per direction used as weights to reflect estimation reliability. Let **R** (**t**) be the 7×1 vector of observed firing rates at time *t*, and let the 7×3 design matrix be

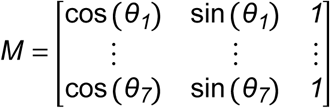

where *θ_1_*, …,*θ_7_* are the seven target directions. The weight matrix *W* = *diag* (*w_1_*, …,*w_7_*) is a diagonal matrix whose entries *w_i_* correspond to the number of trials for direction *θ_i_*. The weighted least squares parameter estimate is

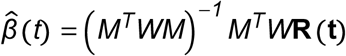

The fitted firing rate for any direction *θ* is obtained as:

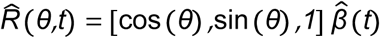

This approach allowed us to estimate SS firing rates for all saccade directions, including those with few or no trials, by leveraging the smooth directional tuning of SS firing rates.

To quantify how much the adaptation-induced changes in SS firing was transferred to non-adapted directions, first, effect size from the baseline to generalization phase was calculated and fitted by a cosine function

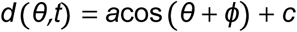

where *ϕ* is the phase shift indicating the direction of peak modulation change relative to the horizontal axis. The in-phase component *IP* = *a*cos (*ϕ*) captures the projection of the cosine fit onto the horizontal axis, reflecting both the magnitude of the adaptation-induced change and its directional alignment with the adapted horizontal direction. This value was compared between CS-on opposed saccades alone and the CS-on opposed minus CS- on aligned difference, to assess whether combining the two subpopulations amplifies the directional signal.

To assess statistical significance, a bootstrap procedure was used. For CS-on opposed and CS-on aligned saccade separately, PC units were randomly resampled with replacement 100,000 times, and the effect size between the baseline and generalization phases was computed for each resample. The in-phase component was then calculated for each bootstrap sample, generating a null distribution against which the observed in-phase component was compared.

### Quantification of CS directional selectivity

The directional selectivity of CS firing around saccade onset was quantified using Cramér’s V, a measure of association between two categorical variables derived from the chi-squared statistic. For each time bin, a 2 × 7 contingency table was constructed, with rows representing CS firing (present or absent within the bin) and columns representing the seven possible target locations. Cramér’s V was then computed as:

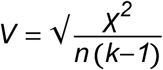

where *χ^2^* is the chi-squared statistic, *n* is the total number of observations, and *k* is the number of categories of the variable with fewer categories (here *k* = *2*). Cramér’s V ranges from 0 to 1, where 0 indicates no association between CS firing and target location and 1 indicates a perfect association.

To ensure sufficient spike counts for reliable estimation, Cramér’s V at each time point *i* was coputed by accumulating CS occurrences within a centered window of ±50 ms, yielding a total window of 101 ms at 1 ms time resolution.

### Effect of peri-saccade onset CSs on saccade kinematics

To assess the effect of peri-saccade onset CSs on saccade kinematics, linear mixed models were used with kinematic parameters (PV, duration, amplitude, and saccade onset position) as dependent variables. For the fatigue paradigm:

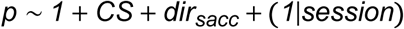

where *p* is a kinematic parameter of interest, *CS* is a binary variable indicating CS presence or absence during the peri-saccade onset period, *dir_sacc_* denotes saccade direction, (CF or CP), and *session* is a random intercept accounting for between-session variability.

For the adaptation paradigm, adaptation direction *dir_adapt_*, and 3 adaptation phases *phase* were included;

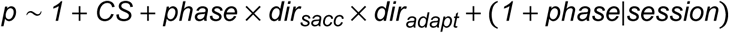

All models were fitted using maximum likelihood estimation (MixedModels.j., Julia). The effects of CS presence on each kinematic parameter was assessed from the fixed effect coefficient of *CS* and its associated Wald z-statistics.

### Assessment of CS firing rate sensitivity to eye position

To assess the onset-position encoding of CS during the peri-saccade onset period, a bootstrap procedure was used. CS-on direction labels were randomly permuted across cells and the slope of the linear regression between CS firing rate and eye position at saccade onset was recomputed for each permutation, repeated 100,000 times. Absolute slope values were used to construct a one-sided null distribution, against which the observed absolute slope was compared. The observed absolute slope was considered significant if it exceeded the 95th percentile of the null distribution (one-sided, *α* =0.05).

Similarly, to assess the difference in slopes between the pre- and post-saccadic error periods for CS-on opposed and aligned saccades, CS-on direction labels were randomly permuted across cells and the slope difference between the two subpopulations was recomputed for each permutation, repeated 100,000 times. This generated a null distribution of slope differences centered around zero. The 95% confidence interval was defined as the 2.5th and 97.5th percentile of this null distribution, and the observed slope difference was considered significant if it fell outside this interval (two-sided, *α* =0.05).

### Comparing SPI across conditions

To test whether the peri-saccadic SPI modulation differed across conditions, we fitted a linear mixed model to the SPI increase from the baseline to the mean time of saccade offset. The baseline was defined the period from -250 to -150 ms relative to saccade onset. The mean saccade offset time was computed separately for each condition. The model had the form

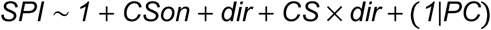

where *CSon* indicates whether the saccade direction was CS-on aligned or CS-on opposed, *dir* indicates CF or CP saccades, and *PC* is a random intercept for each PC. Each PC was tested in only one CF direction, and therefore contributed to one CF condition and to the CP condition in the opposite direction (e.g., CF CS-on opposed and CP CS-on aligned saccades). The random intercept accounts for this repeated sampling. Each observation was the SPI increase of one PC in one condition. Models were fitted separately for the adaptation and fatigue paradigms by maximum likelihood (MixedModels.jl, Julia). Effects were assessed using Wald z-statistics on the fixed-effect coefficients.

### Computing correlation between SPI and CV

Because SPI of individual neurons are too noisy to reliably assess the correlation, we randomly selected 50 cells and computed their mean SPI and CV during -100 to 200 ms from saccade onset across this subset to obtain the Pearson’s correlation coefficient. This sampling procedure was repeated 100,000 times, and the mean correlation and its 95% confidence interval were derived from the resulting distribution.

## Supplementary figures

**Supplementary Figure 1:**
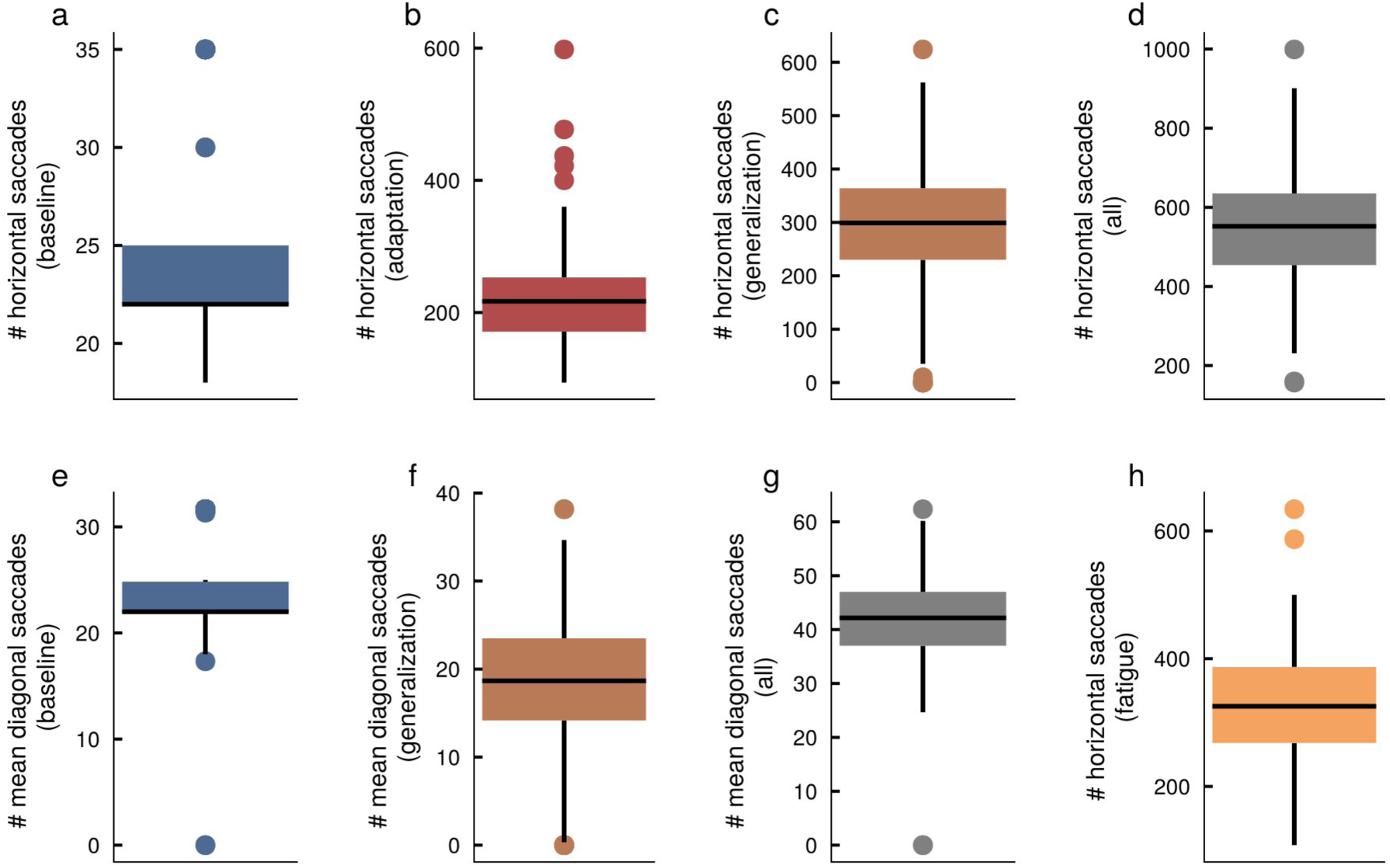
Boxplots of number of trials. **a**-**d**: number of horizontal saccades during the baseline phase (a), the adaptation phase (b), the generalization phase (c), and across the entire session (d). **e**-**g**: mean number of diagonal saccades during the baseline phase (e), the generalization phase (f) and across the entire session (g). **h**: number of saccades recorded in the fatigue paradigm

**Supplementary Figure 2:**
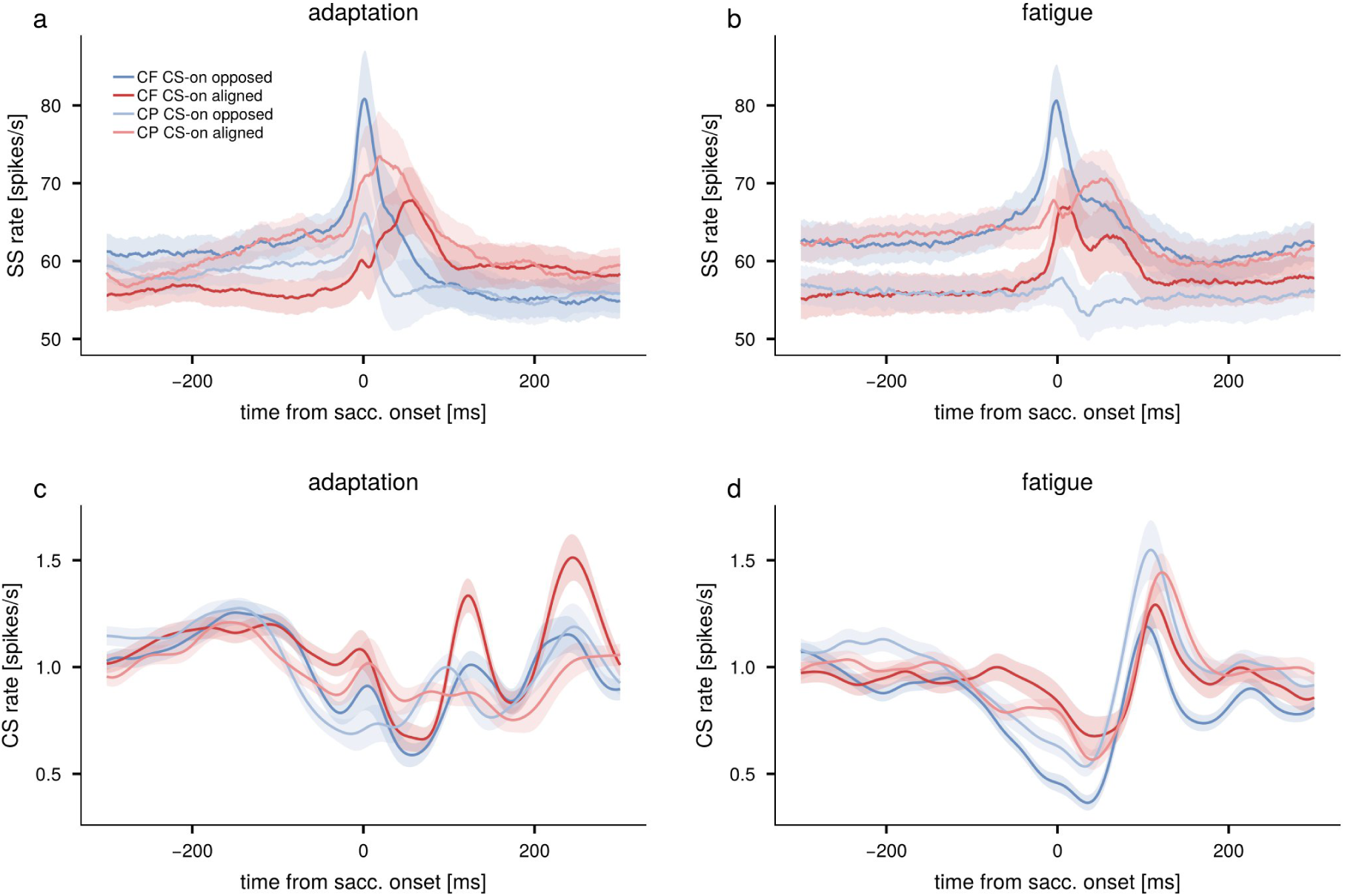
Population responses of SS (**a**,**b**) and CS (**c**,**d**) computed separately for CF (dark shades) and CP (light shades) saccades, shown for CS-on opposed (blue) and CS-on aligned (red) saccades (a,c: adaptation paradigm, b,d: fatigue paradigm). Shaded areas indicate standard error. For the adaptation paradigm, SS population responses were calculated from the baseline phase, whereas CS responses included the entire session to increase the statistical power given the low CS firing rates.

**Supplementary Figure 3:**
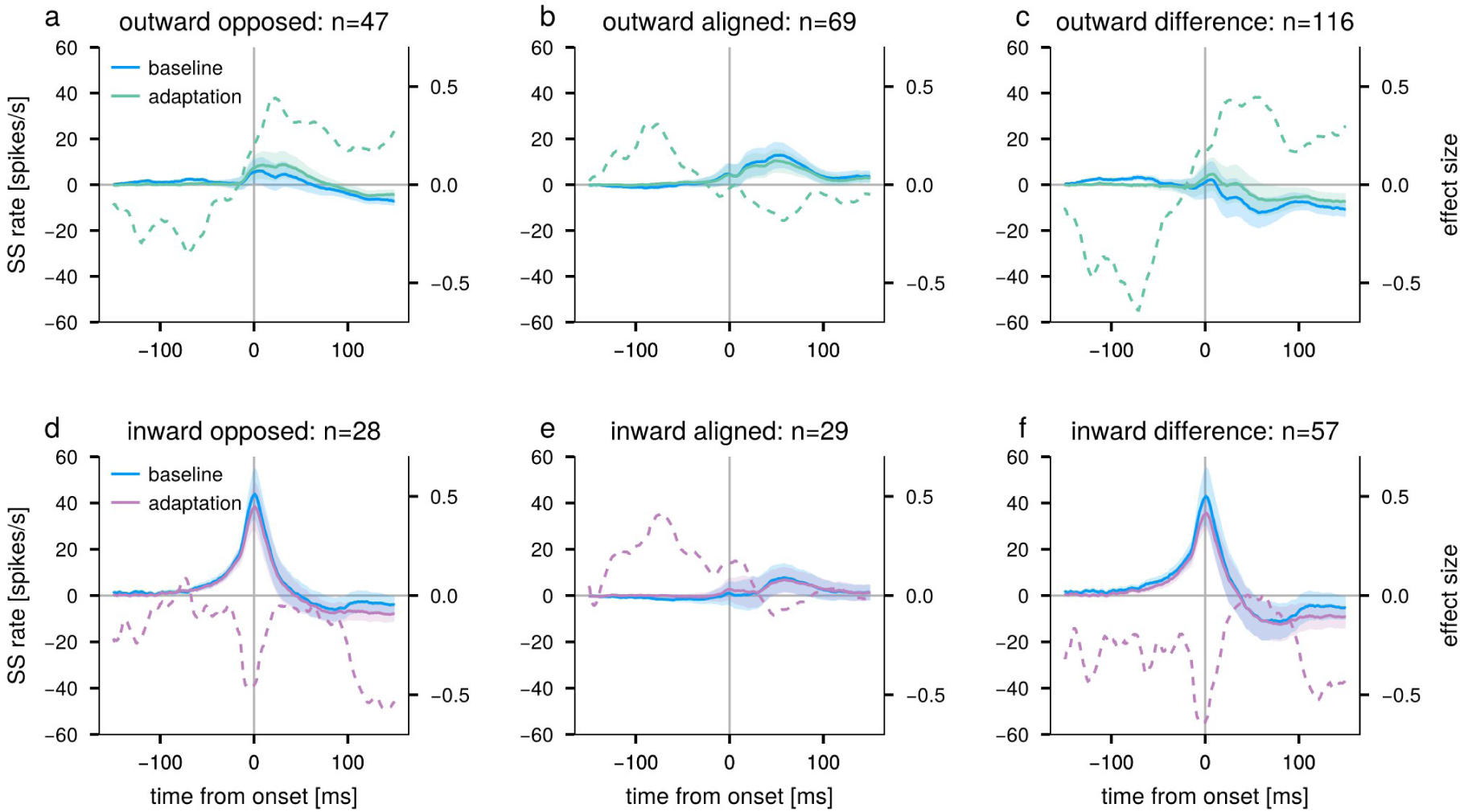
Adaptation-induced changes in SS population firing rate for CS-on opposed (a,d), CS-on aligned (b,e), and their difference (c,f) saccades, shown separately for outward (a-c) and inward (d-f) adaptation. Solid lines show mean SS firing rates during the baseline (blue) and adaptation (green for outward, purple for inward) phases. Dashed lines indicate the effect size of the adaptation-induced change relative to baseline. Shaded areas indicate standard error. The number of PC units contributing each subpopulation is indicated in the panel title.

**Supplementary Figure 4:**
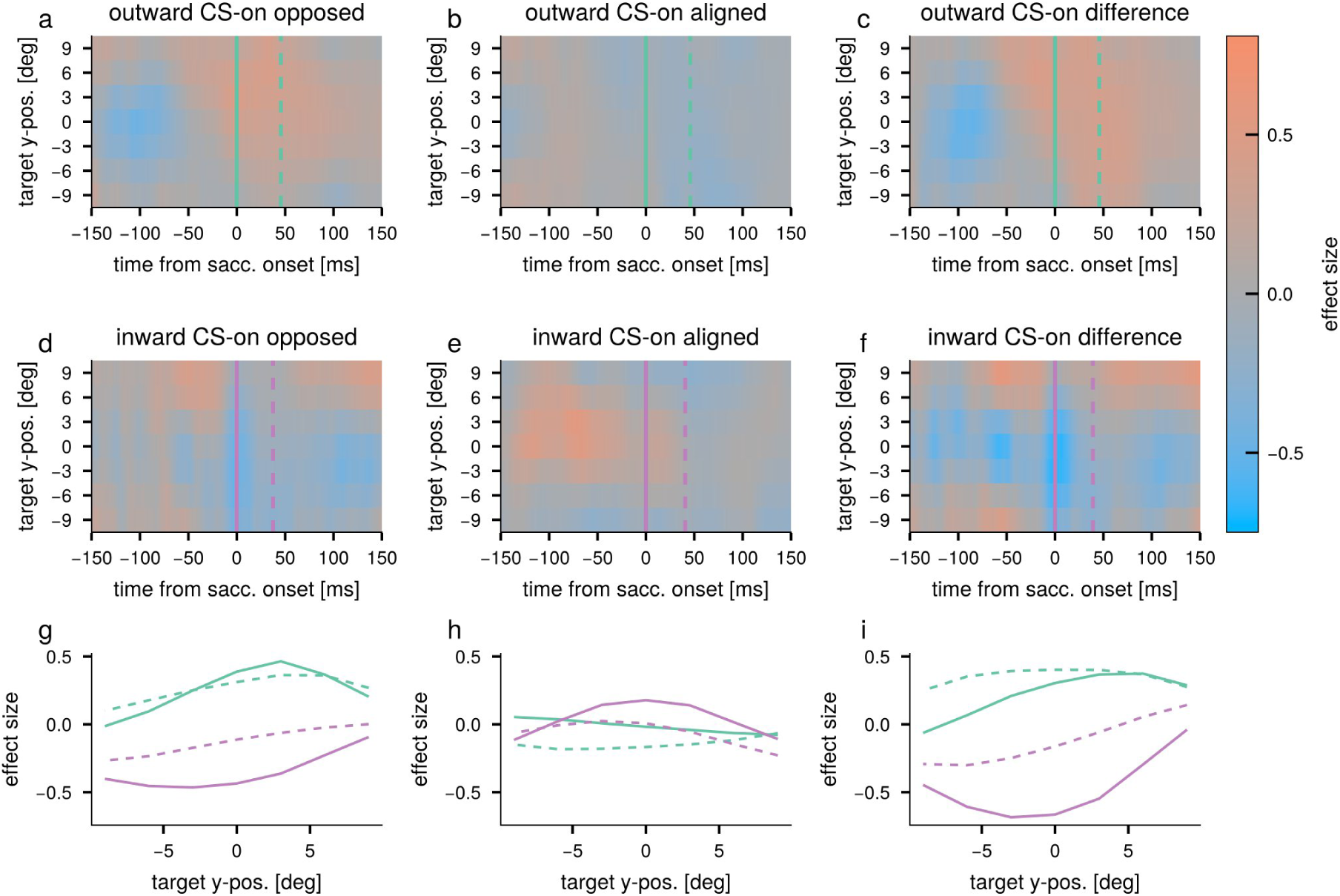
Changes in SS firing after adaptation. **a–c**: Changes in mean SS population response between the generalization and baseline phases of the outward adaptation for CS-on opposed saccades (a), CS-on aligned saccades (b), and their difference (c). Solid and dashed vertical lines indicate saccade onset and mean saccade offset time. **d–f**: Same as a–c, but for the inward adaptation. **g-i**: Changes in mean SS population response after adaptation at saccade onset and mean saccade offset as a function of target position. Line styles and colors match a-f.

**Supplementary Figure 5:**
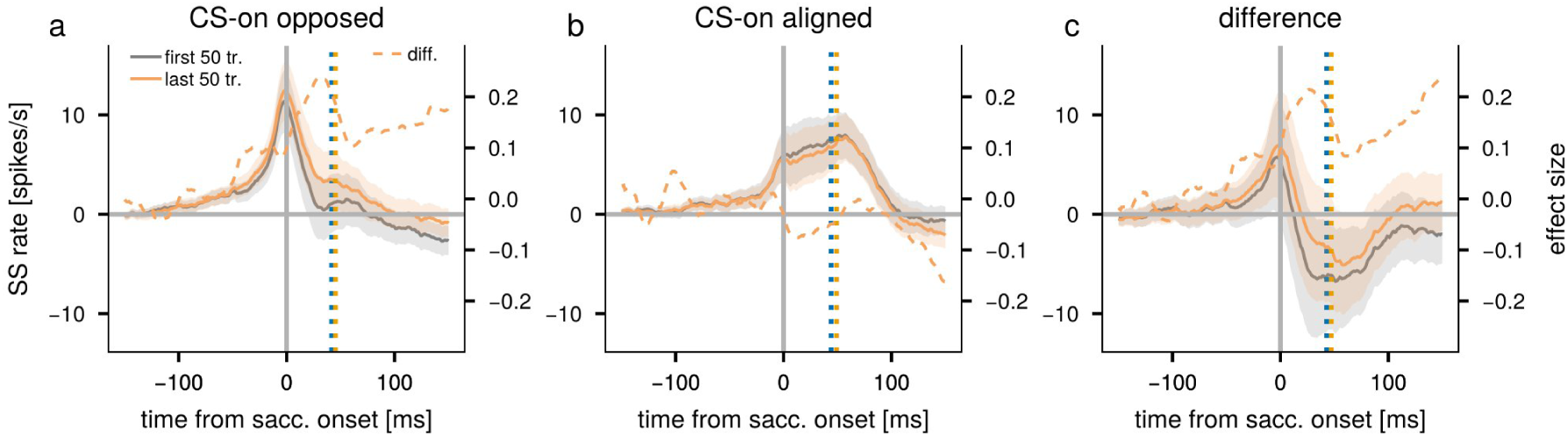
Changes in SS firing after fatigue. **a–c**: Mean SS population responses of the first (gray) and last (orange) 50 trials from the fatigue paradigm, aligned to saccade onset, for CS-on opposed saccades (a), CS-on aligned saccades (b), and their difference (c). Dotted lines indicate the difference between first and last 50 trails as effect size.

**Supplementary Figure 6:**
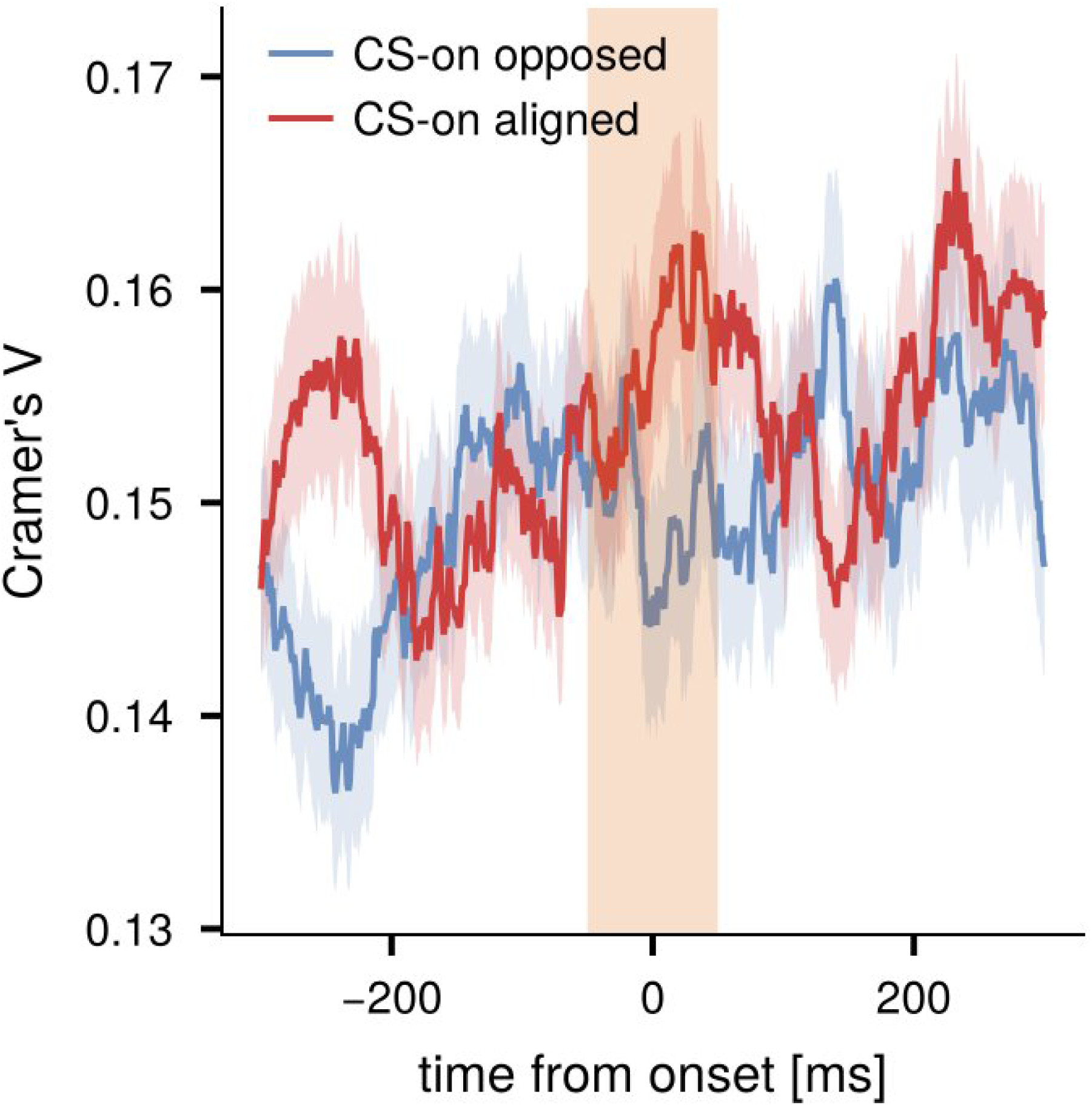
Mean Cramer’s V. Same as Fig. 4 f but for CP saccades.

**Supplementary Figure 7:**
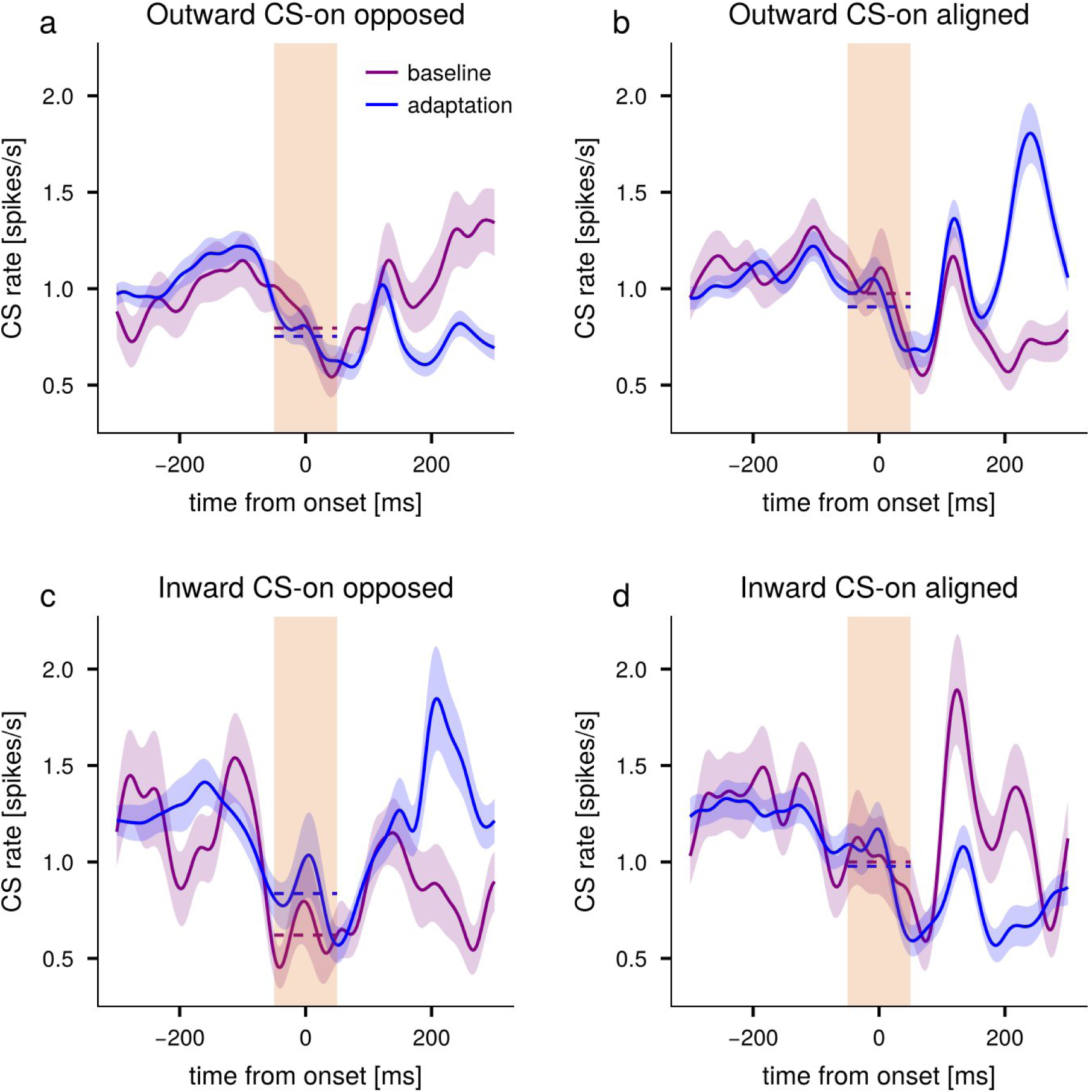
Comparison of CS firing rates between the baseline and adaptation phases for the CF saccades. **a**: CS-on opposed saccades from outward-adaptation sessions. **b**: CS-on aligned saccades from outward-adaptation sessions. **c**: CS-on opposed saccades from inward-adaptation sessions. **d**: CS-on aligned saccades from inward-adaptation sessions.

**Supplementary Figure 8:**
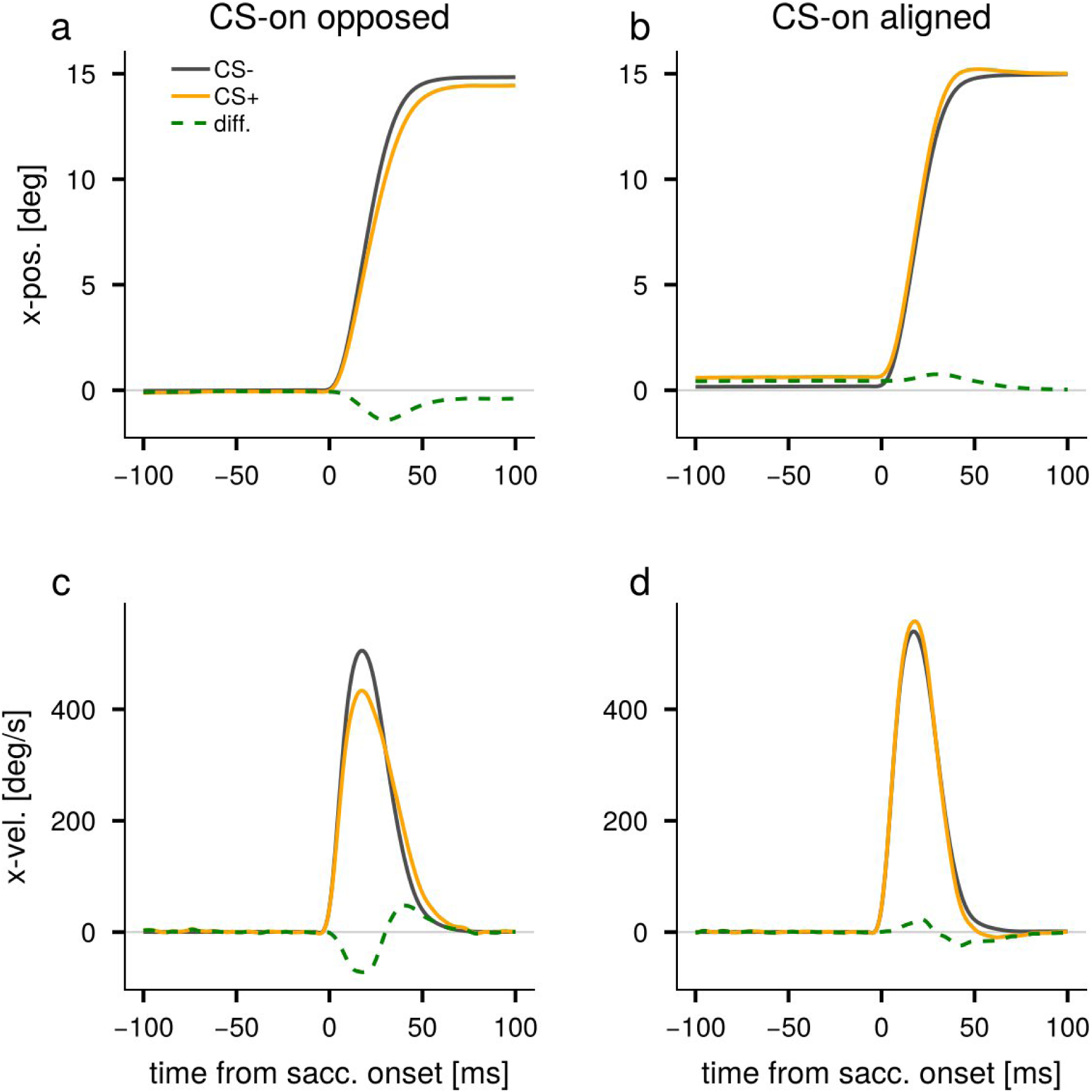
Effect of CS around saccade onset on saccade kinematics in the fatigue paradigm. Same as Fig. 5, but for the fatigue paradigm.

**Supplementary Figure 9:**
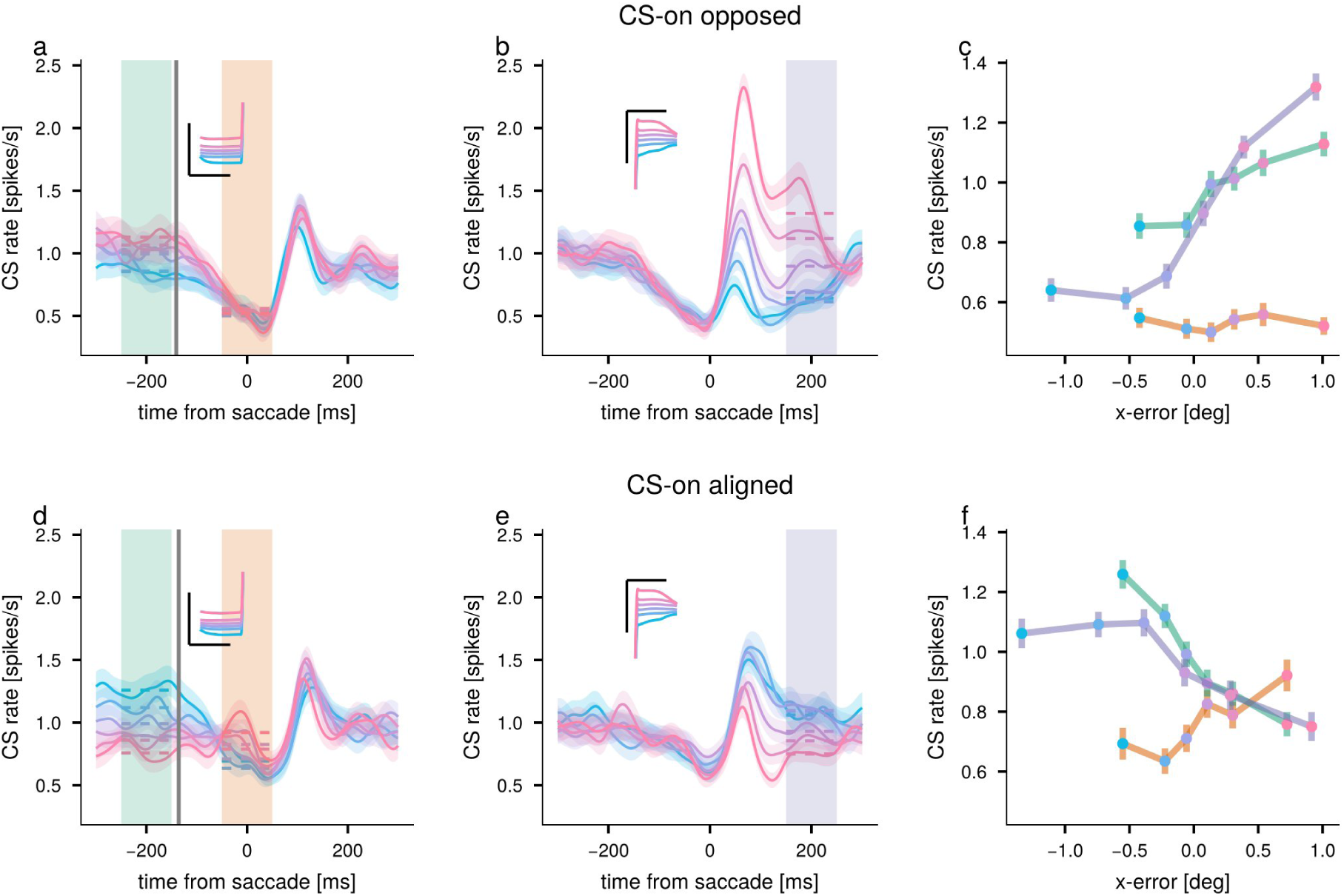
CS firing rate sorted by retinal error in the fatigue paradigm. Same as Fig. 6, but for the fatigue paradigm. Inset: corresponding horizontal eye-position traces; scale bars indicate 3° and 300 ms.

**Supplementary Figure 10:**
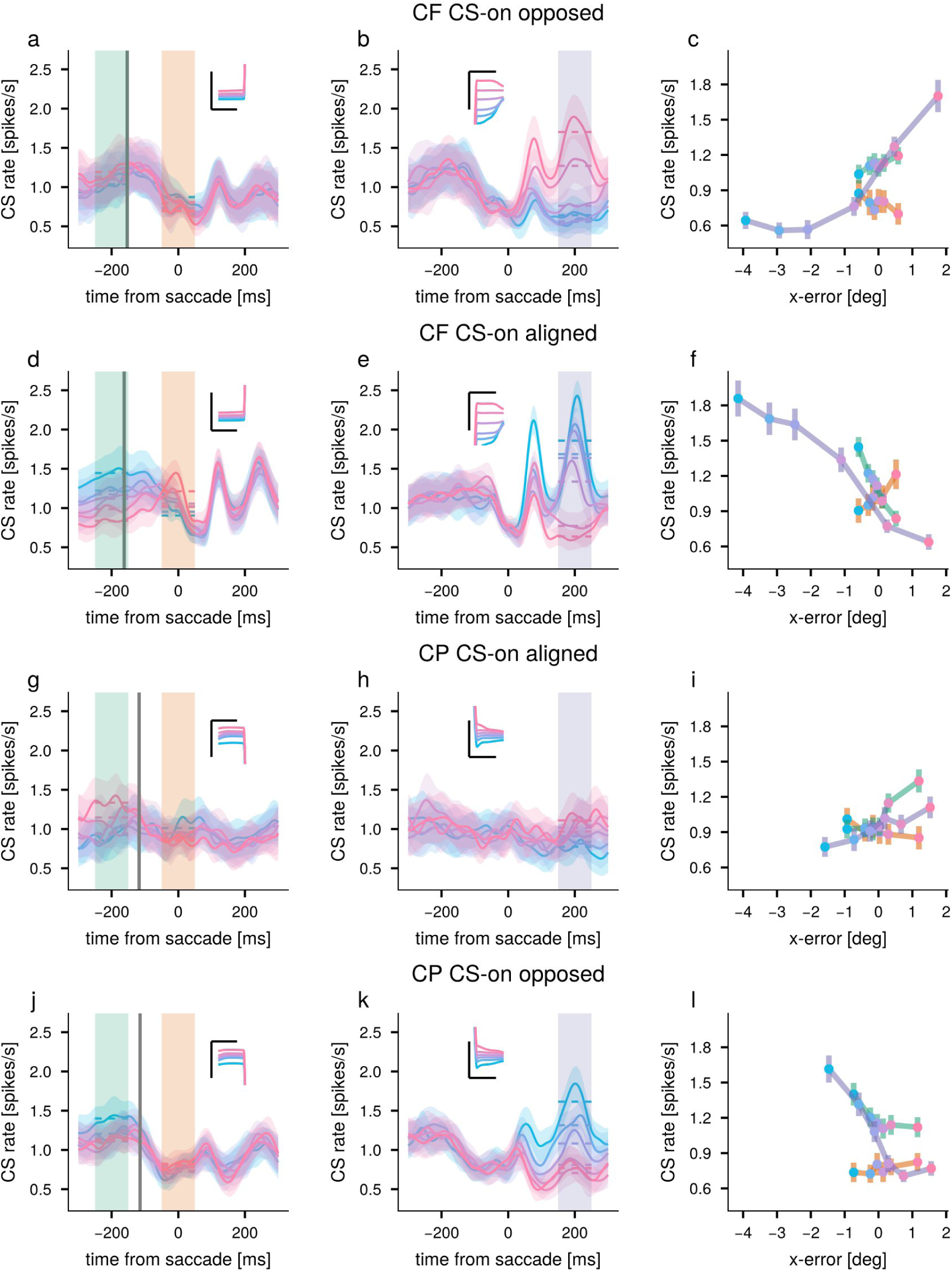
CS firing rate sorted by retinal error during the inward-adaptation sessions. Same as Fig. 6, but restricted to the PCs recorded during outward-adaptation sessions, with CF and CP saccades plotted separately.

**Supplementary Figure 11:**
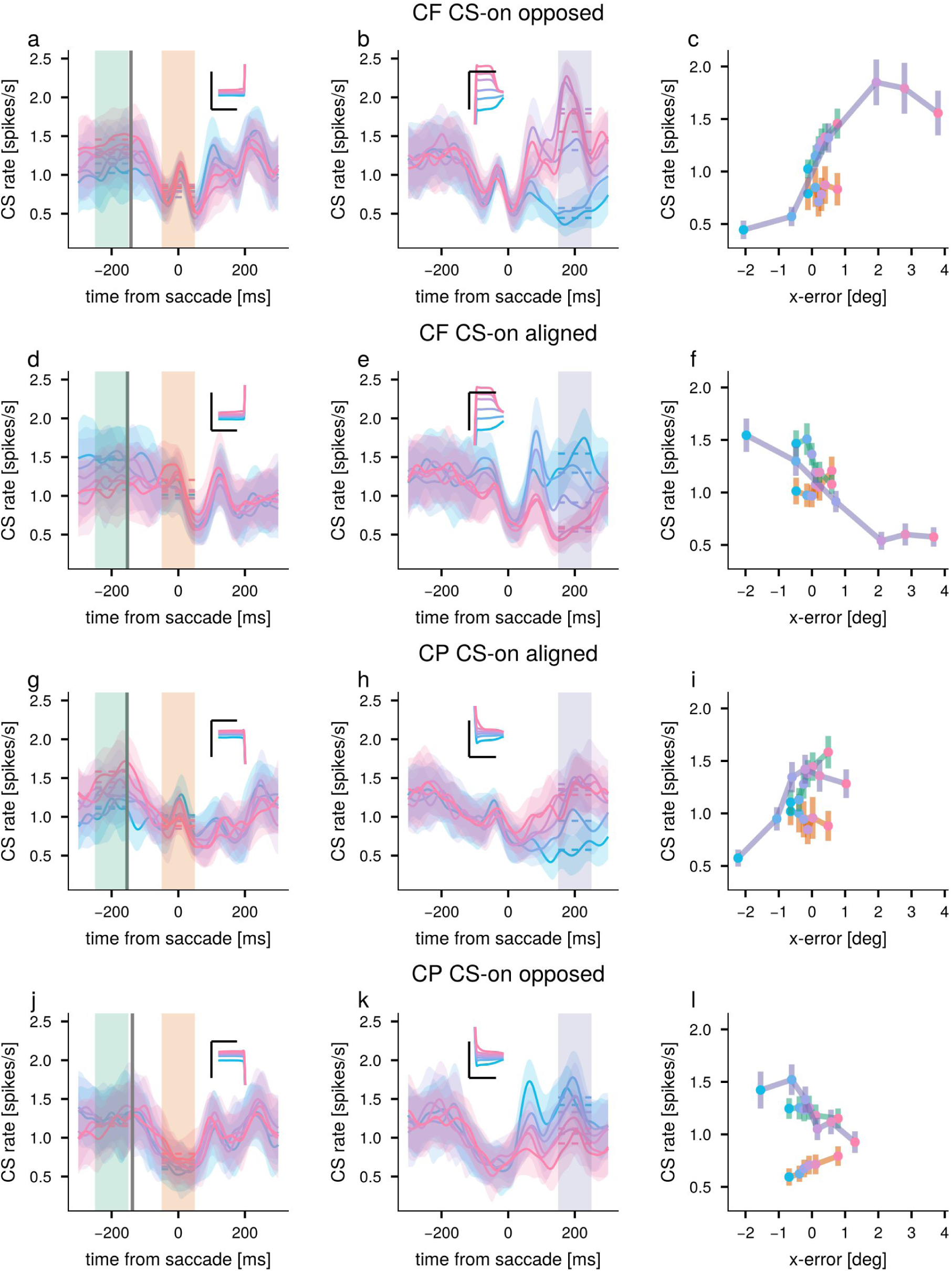
CS firing rate sorted by retinal error during the inward-adaptation sessions. Same as Fig. 6, but restricted to the PCs recorded during inward-adaptation sessions, with CF and CP saccades plotted separately.

**Supplementary Figure 12:**
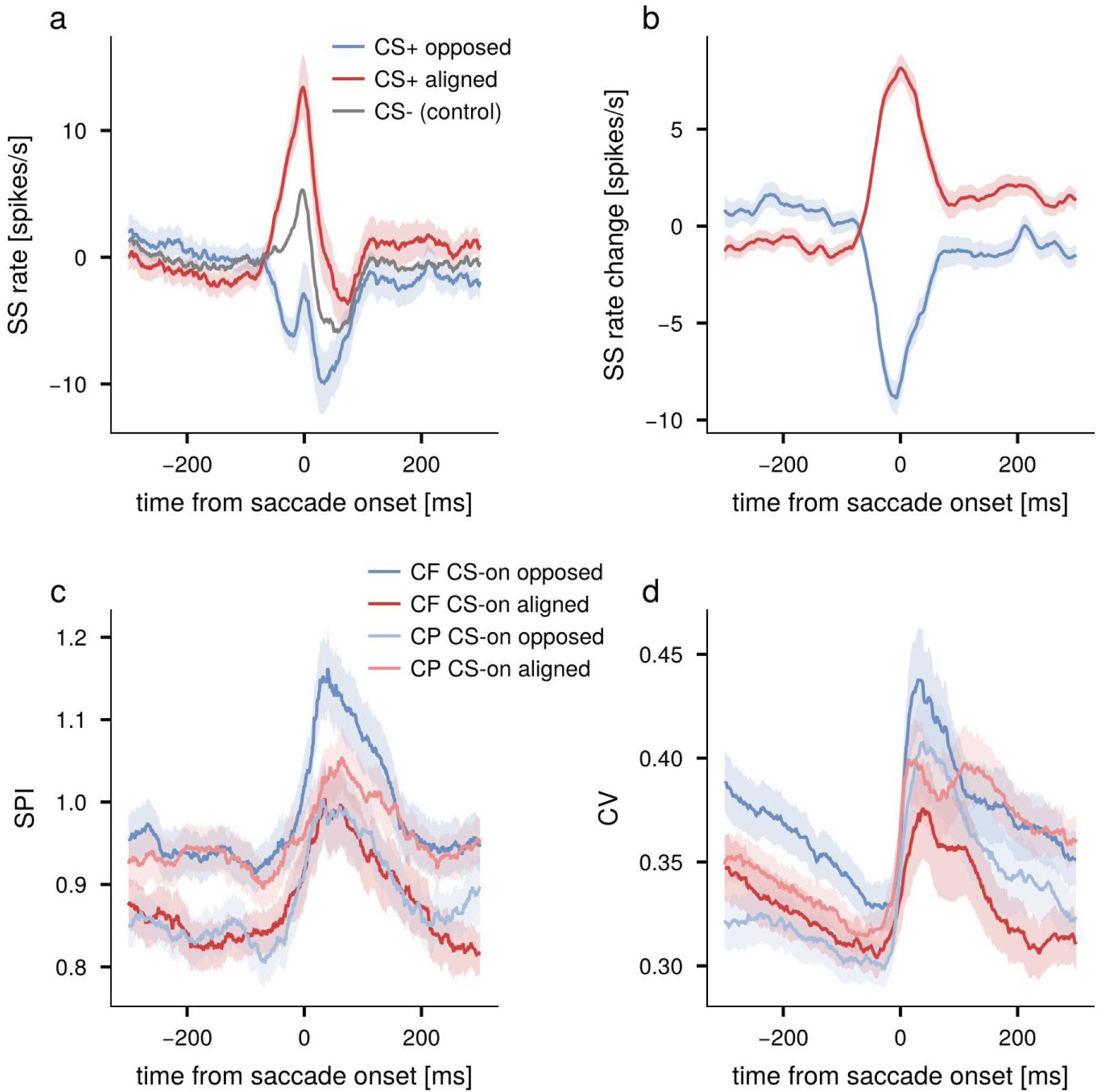
Effect of CS on net SS population responses in the adaptation paradigm. Same as Fig. 7 b-e. but for the adaptation paradigm.

**Supplementary Figure 13:**
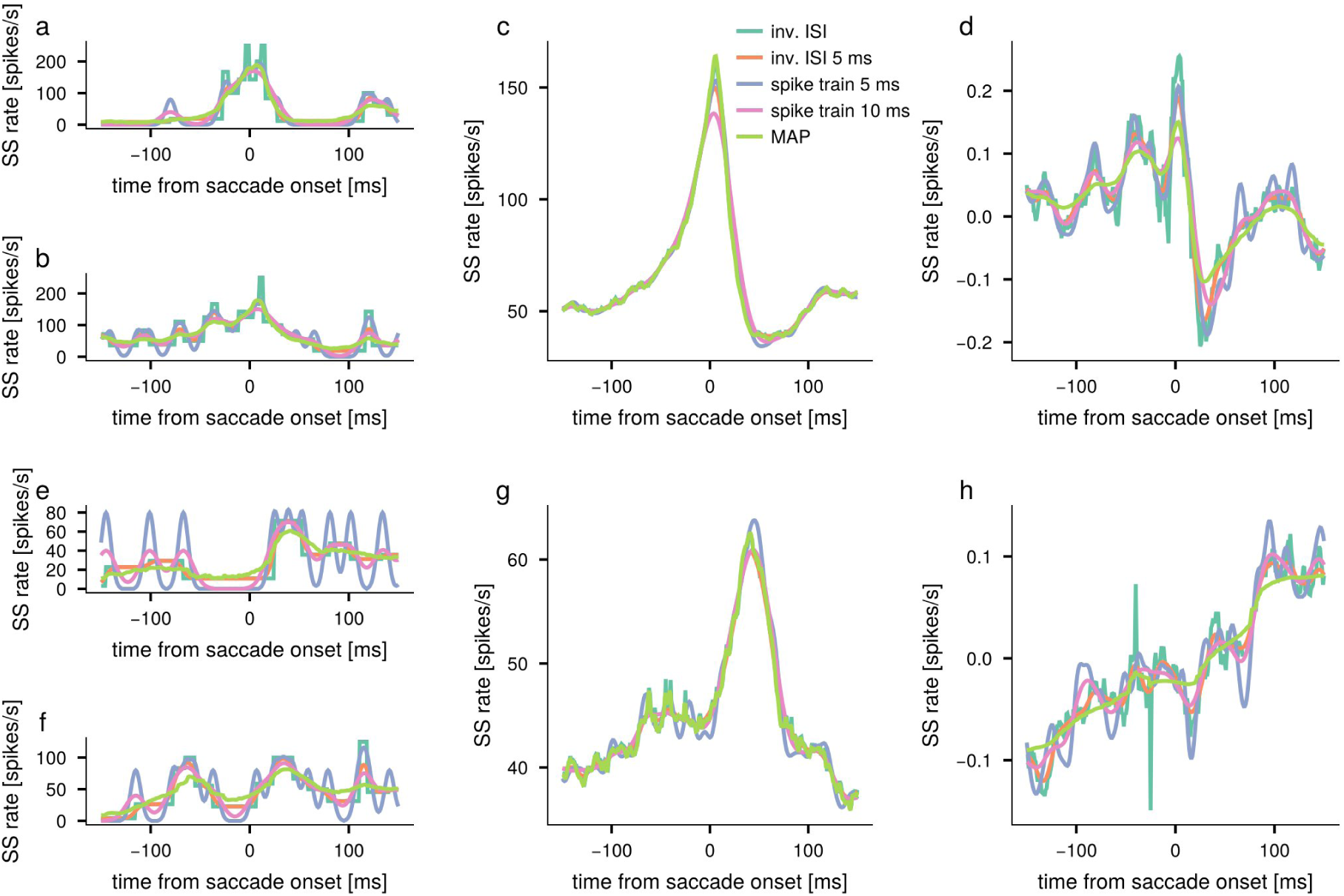
Comparison of estimation methods for SS firing rate for two exemplary PCs. **a,b,e,f**: Instantaneous firing rates from two example trials for the five estimation methods (gray: raw IISI, blue: IISI with a 5 ms kernel, orange: spike train with a 5 ms kernel, green: spike train with a 5 ms kernel, red: MAP). c,g: Mean firing rate. **d,h**: PV sensitivity.

